# *Pseudomonas aeruginosa-*mediated cardiac dysfunction is driven by extracellular vesicles released during infection

**DOI:** 10.1101/2024.11.22.624948

**Authors:** Naresh Kumar, Sameer Salam Mattoo, Shridhar Sanghvi, Maneeth P. Ellendula, Sahil Mahajan, Clara Planner, Joseph S. Bednash, Mahmood Khan, Latha P. Ganesan, Harpreet Singh, William P. Lafuse, Daniel J. Wozniak, Murugesan V.S. Rajaram

## Abstract

*Pseudomonas aeruginosa (P.a.)* is a gram-negative, opportunistic bacterium abundantly present in the environment. Often *P.a.* infections cause severe pneumonia, if left untreated. Surprisingly, up to 30% of patients admitted to the hospital for community-acquired pneumonia develop adverse cardiovascular complications such as myocardial infarction, arrhythmia, left ventricular dysfunction, and heart failure. However, the underlying mechanism of infection-mediated cardiac dysfunction is not yet known. Recently, we demonstrated that *P.a.* infection of the lungs led to severe cardiac electrical abnormalities and left ventricular dysfunction with limited *P.a.* dissemination to the heart tissue. To understand the mechanism of cardiac dysfunction during *P.a.* infection, we utilized both *in vitro* and *in vivo* models. Our results revealed that inflammatory cytokines contribute but are not solely responsible for severe contractile dysfunction in human induced pluripotent stem cell-derived cardiomyocytes (hiPSC-CMs). Instead, exposure of hiPSC-CMs with conditioned media from *P.a.* infected human monocyte-derived macrophages (hMDMs) was sufficient to cause severe contractile dysfunction and arrhythmia in hiPSC-CMs. Specifically, exosomes released from infected hMDMs and bacterial outer membrane vesicles (OMVs) are the major drivers of cardiomyocyte contractile dysfunction. By using LC-MS/MS, we identified bacterial proteins, including toxins that are packaged in the exosomes and OMVs, which are responsible for contractile dysfunction. Furthermore, we demonstrated that systemic delivery of bacterial OMVs to mice caused severe cardiac dysfunction, mimicking the natural bacterial infection. In summary, we conclude that OMVs released during infection enter circulation and drive cardiac dysfunction.

## Introduction

*Pseudomonas aeruginosa (P.a.)* is a gram-negative opportunistic bacterium present in the environment and hospital settings^1,2^. Though *P.a.* infections can occur anywhere in the body, lung infections are often serious and can develop into life-threatening diseases with increased mortality^3^. In the US, over 5 million people develop pneumonia annually, ranking as the 8th leading cause of death^4^. It is reported that up to 30% of patients hospitalized with community-acquired pneumonia (CAP) develop cardiovascular complications^5^. Older adults with CAP are more likely to have comorbidity with cardiovascular diseases (CVD) such as ischemic heart failure, diabetes, stroke, and chronic obstructive pulmonary disease (COPD)^6^. Several research studies have been published linking pneumonia with CVD^3,6–9^, and recently, we demonstrated that *P.a.* infection causes cardiac dysfunction with low to minimal *P.a.* dissemination into the heart tissue^10^. However, the mechanism of CVD during *P.a.* infection has not yet been clearly studied. We hypothesize that the release of effector molecules, such as exosomes from infected cells and outer membrane vesicles (OMVs) from bacteria, travelled from the lungs and caused cardiovascular complications.

Once *P.a.* infection is established in the lungs (in the case of pneumonia) or at other infection sites, the bacteria release bacterial pathogen-associated molecular patterns (PAMPs) and OMVs. The infected cells release inflammatory mediators such as IL-1β, TNF-α, IL-6, and damage-associated molecular patterns (DAMPs) into the bloodstream which activate nonspecific inflammation and tissue damage. It has been known that the Gram-negative bacteria secrete nano-size OMVs packed with several immunogenic molecules, including endotoxins, enzymes, peptidoglycan, periplasmic proteins, short RNAs (sRNAs), and nucleic acids^11–14^. The size of the OMVs varies from 10-300 nm^15–19^ in diameter, and they play a vital role in host-pathogen interactions where they mediate intracellular communication, modulate the host immune response, and function in distal organs^11–14^. Like the OMVs released by bacteria, host cells also release extracellular vesicles (EVs). Based on the size and mechanism of release, these vesicles have been classified into three types. EVs larger than 1000 nm are called microvesicles, those less than 1000 nm are called apoptotic bodies, and EVs with an average size of 40-160 nm are called exosomes^20–23^. Earlier studies demonstrated that the process of exosome secretion is a way to release cellular waste^20,24,25^. Exosomes carry important proteins and genetic materials which have been shown to play a vital role in cell-cell communication and have also been associated with physiological functions and pathogenesis.

Recently, hiPSC-CMs have gained significant attention as a powerful *in vitro* model system to study the mechanisms underlying cardiomyopathies, cardiotoxicity, and are used for drug screening^26–28^. Moreover, advancements in hiPSC generation techniques, the scalability of hiPSC-CMs, and the emergence of next-generation genetic manipulation methods make hiPSC-CMs a promising model for developing patient-specific precision medicine^28,29^. Furthermore, hiPSC-CMs express similar cardiac ion channels as mature adult cardiomyocytes of heart tissue and have a similar action potential duration, comparable to the human QT interval^30,31^. Therefore, in this study, we used hiPSC-CMs as a platform for testing cardiomyocyte contractile function during bacterial infection and dysfunction induced by bacterial-derived components.

In the current study, we treated hiPSC-CMs with conditioned media (C-media) harvested from *P.a.* infected hMDMs, or OMVs from *P.a.,* and studied cardiomyocyte contractile function. We found that exposure of hiPSC-CMs to C-media from hMDMs infected with *P.a.* caused severe contractile dysfunction with a decreased and irregular beat period, decreased field potential duration (FPD), and slower conduction. Furthermore, C-media exposure also caused arrhythmia which were confirmed by prolonged action potentials with Early after Depolarization (EAD) shoulders, increased Action Potential Durations (APDs), triangulation ratio, and rise time. Furthermore, we found similar contractile dysfunction when hiPSC-CM were exposed to purified bacterial OMVs. Finally, after confirming our *in vitro* findings, we performed *in vivo* mouse experiment, in which we injected OMVs and monitored the mice survival and heart function. Our data revealed that the OMVs caused severe cardiac electrical dysfunction and arrhythmia. In conclusion, the exosomes released from infected cells and OMVs released from *P.a.* carry both host- and bacterial products. OMVs that enter circulation are able to enter the heart and cause cardiac dysfunction.

## Material and methods

### Ethics statement

All animal procedures in this study were approved by the Institutional Biosafety Committee (IBC) and The Ohio State University Institutional Animal Care and Use Committee (IACUC) under protocol 2020A00000004.

### Bacterial growth and media

*P.a.* (strain PAO1) was grown overnight in Luria broth (LB) at 37°C and used as a starter for fresh *P.a.* culture for infection experiments. Briefly, 100 μl of overnight grown *P.a.* were inoculated with 5 mL LB and allowed to grow until log phases (0.5–0.8 OD at 600 nm); the bacteria were harvested and resuspended in RPMI or saline for *in vitro* and *in vivo* experiments, respectively.

### Isolation of Human monocyte-derived macrophages (hMDMs)

Human monocyte-derived macrophages (hMDMs) were prepared from healthy human volunteers using an approved OSU IRB protocol as described^32^. Briefly, PBMCs were isolated from heparinized blood on a Ficoll cushion and then cultured in Teflon wells (Savilexx, Minnetonka, MN, USA) for 5 days in 20% autologous serum. hMDMs in the cultured PBMCs were adhered to tissue culture plates for 2-3h at 37°C and 5% CO2 in 10% autologous serum. Lymphocytes were washed away, and hMDM monolayers were repleted with RPMI containing 10% autologous serum and incubated overnight before being used for infection experiments.

### The culture of human induced pluripotent stem cell-derived cardiomyocytes

hiPSC-CMs were plated according to the supplier’s protocol as described earlier^32^. Briefly, hiPSC-CMs was cultured in 0.1% gelatin coated tissue culture plate using plating media (FUJIFILM Cellular Dynamics, Inc. WI, USA) and incubated for 48 h in a humidified chamber at 37 °C and 5% CO2. Subsequently, we replaced the plating media with CDI maintenance medium (FUJIFILM Cellular Dynamics, Inc. WI, USA) and changed every other day for 5-7 days. hiPSC-CMs from were harvested and re-plated (3.0X10^4^ cells per well) in a 24-well CytoView MEA plate (Axion BioSystem, GA, USA) coated with fibronectin (50 µg/mL) and incubated at 37°C with 5% CO2 in a humidified atmosphere, and the maintenance medium was changed every alternate day and used for our *in vitro* assays.

### Infection of hMDMs and processing of conditioned media (C-media)

Overnight cultured hMDMs were infected with 1 MOI of *P.a.* (with live or heat-killed at 95°C for 45 minutes) for 22h. The cell-free culture supernatants from un-infected and *P.a.* infected plates were harvested, centrifuged at 10,000g for 10 minutes to remove the debris, filter sterilized using a 0.22 µm filter (now called C-media) and stored at -80°C until used for hiPSC-CMs stimulation experiments. To confirm the absence of active bacteria in C-media, a CFU assay was performed using 10 µl of C-media plated on agar plate (PIA medium, BD, MD USA), incubated at 37°C overnight and colonies were counted.

### Treatment of hiPSC-CMs and MEA

hiPSC-CMs was seeded in MEA plate for 7-10 days for synchronization and monolayer formation. Then, hiPSC-CMs were exposed to C-media obtained from live *P.a.* or heat-killed *P.a.* (HI) and/or left untreated (UT) in a ratio of 1:1 (v/v) of hiPSC-CM media and C-media. Real-time cardiomyocyte functionality data was acquired on MEA as previously described^33^. In brief, cells were stabilized for a minimum of 30 min in the MEA system (Maestro Edge, Axion Biosystem, GA, USA), then the baseline was recorded, followed by C-media treatment. The data was recorded for 5 minutes at an interval of 15 minutes for the mentioned duration using AxIS Navigator™ software version 2.0.4.21. The data analysis was performed using Cardiac Analysis tool™ version 3.1.8 (Axion Biosystem, GA, USA) and AxIS metric plotting tool version 2.3.1.

### Cytotoxicity assay

Cell death was assessed by using the Cytotoxicity Detection Kit (Roche, Germany) following the manufacturer’s instructions. Briefly, 25 µL of culture medium was transferred to a 96-well plate, mixed with 25 µL of the LDH cytotoxicity assay reagent, incubated for 30 minutes at 37°C, and the optical density was measured at 492 nm with Spectrophotometer (Molecular Devices).

### ELISA

Cell-free culture supernatants (C-media) from P.a. infected or OMV exposed hMDMs were collected at 24h and used to measure IL-1β, IL-6, TNF, and IL-10 by ELISA (Duo set ELISA kits, R&D Systems) according to the manufacturer’s instructions.

### Isolation and quantification of outer membrane vesicles (OMVs) and Extracellular vesicles (EVs)

*P.a.* (PAO1 strain) was cultured overnight in 10 mL LB. The next day, 500ml LB culture was inoculated with 10mL *P.a.* culture and further grown in a bacterial incubator at 37°C for 20-22h. Then, the bacterial culture was centrifuged at 5,000g for 15 min at 4°C. The supernatant was collected and further centrifuged at 10,000g for 30 min to remove cell debris. The supernatant was collected and passed through a 0.45 µm PES filter to remove the bacteria. Then the OMVs were pelleted by ultra-centrifugation at 150,000g for 2 hours at 4°C and the OMV pellet was washed with PBS. Finally, the OMVs were resuspended in PBS and stored at -80°C for further analysis and experiments. The same protocol was used for the isolation of EVs (mixture of exosomes and OMVs) from the C-media harvested from *P.a.* infected hMDMs. NanoFCM (NanoFCM Co., Ltd, UK) was used for the quantification of OMVs and EVs.

### Separation of OMVs from EVs for LCMS analysis and Western blotting

Extracellular vesicles were harvested from C-media from *P.a.* infected hMDMs and were incubated with CD9 magnetic beads to separate OMVs from EV fraction. For this, 100 µl of human CD9 magnetic beads (Invitrogen) were taken into 1.5ml tubes and washed three times with 1 ml PBS on a magnetic stand. Then, 100 µl of concentrated EVs were mixed with CD9 magnetic beads in a 1:1 ratio and incubated overnight at 4°C. The tubes were placed in a magnetic stand, and the unbound sample was collected (flowthrough), mixed again with fresh CD9 magnetic beads, and incubated overnight at 4°C. The exosome-free vesicles (OMVs) and the CD9-bound exosomes were lysed with TN-1 lysis buffer^34^. Protein quantification was done using the BCA method and lysates (25 µg) were subjected to LC-MS/MS analysis. Protein matched lysates from CD9 bound EVs and free OMVs were subjected to Western blot analysis to confirm the purity of EVs and OMV separation. We probed the membrane with an anti-CD9 antibody followed by a specific secondary antibody and development by use of ECL (Amersham Biosciences/ GE Healthcare). Similar methods were used to determine the flagellin B levels in the exosomes harvested from BALF of *P.a.* infected mice and human serum samples from ICU patients.

### Intracellular Ca^2+^ measurements

For intracellular Ca²⁺measurements, hiPSC-CMs were plated onto fibronectin-coated 29 mm glass-bottom dishes. The Ca²⁺ transients in the hiPSC-CMs were visualized using Fluo-3AM (ThermoFisher Scientific) according to the manufacturer’s protocol and published protocol^33^ with some modifications. Briefly, hiSPC-CMs were incubated with DMEM containing 10µM Fluo-3AM dye for 40 mins at 37°C in a humidified 5% CO2 incubator. Subsequently, the medium was replaced with fresh medium and C-media in a 1:1 ratio, and the cells were further incubated for 30 minutes before imaging. Intracellular Ca²⁺ was concurrently monitored through line scan imaging using a Nikon A1R laser-scanning confocal microscope equipped with a 60× 1.4 NA oil-immersion objective under 488 nm excitation, with emitted light collected in the 500–530 nm range. All experiments were performed at room temperature with triplicate wells.

### Animal experiments

The C57BL/6J mice used were obtained from Jackson Laboratory and housed at The Ohio State University’s Animal Resources Facility. Healthy male and female mice at 10- 12 weeks old were used in this study. Mice were kept on a 12:12h light cycle at 30-70% humidity, with standard water and chow.To minimize the potential confounders, simple ramdomization was performed by including the age matched mice and treating the mice mice with vehicle only. All animal handling and infections were done in an animal facility at Ohio State University and followed OSU’s IACUC guidelines. Mice were injected intravenously with 200 µl DPBS (Mock) or OMVs (final OMV per mice = 10 mg/kg body weight in a 200 µl volume) and were observed for 36h. The blind method was used for group allocation of the mice, and it was masked from the laboratory study team. Standard early removal Criteria, such as >20% weight loss, body score and weakness was considered for removing the mice from the study.

### Electrocardiography (EKG) and Echocardiography (Echo)

For subsurface ECG recordings, 1.5% Isoflurane in oxygen was used as anesthesia at a flow rate of 1.0 L/min. Mice were placed supine on a heated pad to regulate body temperature, and subcutaneous electrodes were placed beneath the skin in a lead II configuration. ECGs were recorded for five minutes each on a PowerLab 4/30 (AD Instruments, Houston, TX). The mice remained unconscious during the reading. Before data analysis, the ECG tracings were manually checked for anomalies or artifacts. The ECG traces were then analyzed using AD Instruments’ LabChart 9 Pro software. To assess cardiac function *in vivo*, 2D-Echo (Vevo 2100, Visualsonics) was performed in mock or OMV-injected mice, 24 hours after administration. Mice were anesthetized in an induction chamber with 1.5% isoflurane in oxygen and a flow rate of 1.0 L/min. Mice were then placed supine on a heated stage, and hair on the chest was removed with depilatory lotion. Isoflurane 1.5% was used to maintain anesthesia. Using an MS-400 transducer, proper anatomical orientation was determined by imaging the heart’s parasternal long axis. M-mode images were taken at the level of papillary muscles. The images were analyzed to determine heart functions.

### Human Serum sample

The Ohio State University ICU Registry and Prospective Cohort Study (BuckICU) (IRB: 2020H0175, IBC: 2020R00000034) is a collaborative effort among investigators within the Division of Pulmonary, Critical Care and Sleep Medicine and Department of Internal Medicine. As co-investigators on BuckICU-SARS, we will have access to 1) blood specimens collected at days 1, 3, 7, 10 and 21; 2) endotracheal aspirates at days 1 and 7; and 3) bronchoalveolar lavage fluid (as clinically indicated), in addition to longitudinal clinical data. This study began collecting samples from adults admitted to the Ohio State University Wexner Medical Center and James Cancer Hospital during the 2020 SARS-CoV-2 COVID-19 pandemic. In January 2022, the study was expanded to include critically ill patients with respiratory failure and/or sepsis, regardless of etiology. This strategy establishes multiple groups for post hoc comparison, including patients with 1) sepsis, 2) septic shock, 3) mild/moderate/severe ARDS, and 4) risk of ARDS or sepsis.

### Statistical Analysis

Statistical analyses were conducted using R software. Mean comparisons were performed using either the Two-tailed Student’s T-test or two-way ANOVA, followed by Tukey’s post hoc tests where applicable. Results are presented as Mean ± SEM, with statistical significance defined at p < 0.05.

## Results

### Conditioned media from *P.a.* infected hMDMs induce cardiomyocyte contractile dysfunction

*P.a.* causes lung infections in humans that can result in severe pneumonia and heart failure^10,35^. We have previously demonstrated that *P.a.* infection in mice causes cardiac electrical and left ventricular dysfunction without any considerable amount of bacterial growth in the heart^10^, which suggests that host-derived inflammatory mediators or bacterial-derived pathogen-associated molecular patterns (PAMPs) released into circulation enhances the systemic inflammation mediated cardiac dysfunction or may enter the heart to induce the cardiac complications. To determine whether *P.a.* infection causes cardiac dysfunction by releasing PAMPs and inflammatory mediators, we infected hMDMs with *P.a.*, collected C-media and filter sterilized to remove live bacteria, and used for *in vitro* experiments. We confirmed that the C-media harvested from *P.a.* infected hMDMs did not contain live bacteria. hiPSC-CMs cultured on a Multielectrode Array (MEA) plate were then exposed to the C-media. The electrical activity of the hiPSC-CMs was monitored over a period of 24 h, and the acquired data was analyzed using the AxIS navigator software. The MEA data revealed that hiPSC-CMs treated with C-media harvested from *P.a.* infected hMDMs significantly inhibited their electrical activity (**Fig. 1A**), as evidenced by a shorter beat period at 60 min compared to the hiPSC-CMs treated with C-media from uninfected hMDMs (**Fig. 1B**). Also, we found that exposure of C-media harvested from *P.a.* infected hMDMs significantly increased the beat rate of hiPSC-CMs at 45 min and then declined with contractions stopping at 120 min post-treatment (**Fig. 1C**). Interestingly, we observed that the C-media-treated hiPSC-CMs appeared to have normal morphology 3 hr post-treatment and maintained viability up to 24 hr. post-treatment (**S.Fig. 1A and B)**. Next, we measured FPD, the time interval between depolarization and repolarization of cardiomyocytes. hiPSC-CMs treated with C-media harvested from *P.a.* infected hMDMs significantly reduced FPD at 60 and 120 min (**Fig. 1D**). Furthermore, the conduction plot shown in **Fig. 1E** represents the synchronized beat of cardiomyocytes. The blue color represents faster wave propagation (faster cell-to-cell communication in a shorter time) while the red color represents a slower wave propagation, which indicates that treatment of hiPSC-CMs with C-media from *P.a.* infected hMDMs disrupts the syncytium of cardiomyocytes and delayed wave propagation. Additionally, treatment of hiPSC-CM with *P.a.* C-media accelerates the prognosis of arrhythmia as evidenced by irregular beat periods and missed depolarization spikes **(Fig. 1F-I**). Together, these data indicate that C-media from *P.a.* infected hMDMs induces cardiomyocyte contractile dysfunction as demonstrated by decreased beat rate, FPD, abnormal conduction, and irregular beat period (abnormal automaticity), and indicated that it could be associated with a proarrhythmic condition.

**Figure 1:**
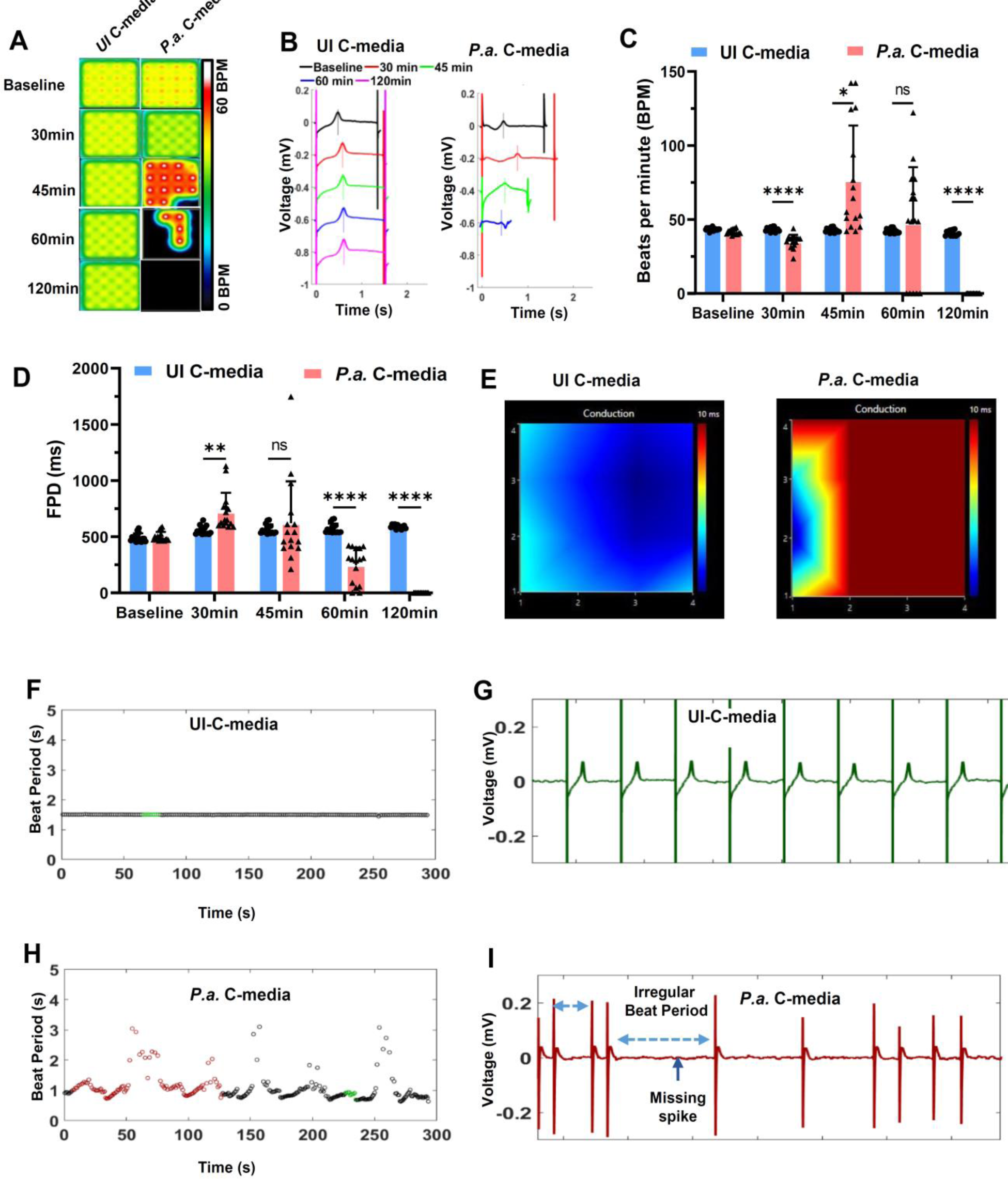
C-media from *P.a.* infected hMDMs cause cardiomyocyte contractile dysfunction. hiPSC-CMs were plated in 24 well MEA plates, and the cells were exposed to the mixture of cardiomyocyte culture medium and C-media (1:1 ratio) harvested from uninfected (UI C-media) or *P.a.* infected (*P.a.* C-media) hMDMs. The cardiomyocyte contractility and electrical activity were recorded using AxIS Navigator on the MEA system at 5% CO2 and at 37°C for indicated time points. Data analysis was performed using the cardiac analysis tool. (**A**) Electrical activity map showing the changes in the beat rate of cardiomyocytes treated with UI C-media and *P.a.* C-media. The activity map shown is a representative well from quadruplicate samples for each treatment and four repeats (N=4). (**B**) Representative traces recorded with the MEA system showing the beat period, T-wave, and FPD in UI-C-media and *P.a.* C-media-treated hiPSC-CMs. The graphs shown in (**C**) are beats per minute and (**D**) is filed potential duration (FPD) at baseline, 30-,45-, 60- and 120 minutes post treatment of hiPSC-CMs with UI C-media and *P.a.* C- media. (**E)** Representative wave propagation of hiPSC-CMs treated with UI C-media and *P.a.* C-media. (**F-G)** Representative EKG traces of hiPSC-CMs treated with UI C-media (**H-I**) *P.a.* C-media. Data shown in Figures C and D are accumulative data from 4 independent experiments, mean ± SD: * p<0.05, ** p <0.01, *** p < 0.001, **** p < 0.0001.

### Conditioned media from *P.a.* infected hMDMs induces arrhythmia in hiPSC-CMs

An irregular beat period suggests the prognosis of arrhythmia, which can be identified by measuring the action potential (AP)^36,37^. In normal physiological conditions, the inward current of Na^+^ and Ca^2+^ is counter balanced by the outward K^+^ current for a specific time, and this follows a specific pattern. The variation in local extracellular action potential (LEAP) in cardiomyocytes exposed to C-media was determined using MEA system. We treated hiPSC-CMs with *P.a.* infected and control C-media and recorded physiological parameters on MEA at different time points. *P.a*. C-media treatment of hiPSC-CMs prolonged the AP with EAD shoulders at different time points (**Fig. 2A**), and 80% of the beats showed EAD at both the 30- and 45-min treatment times (**Fig. 2B**). Furthermore, we compared the Action potential duration (APD) at -30 %, -50%, and 90% repolarization of hiPSC-CMs treated with *P.a.* C-media and found a significant prolongation of AP compared to cardiomyocytes treated with C-media from uninfected hMDMs (**Fig. 2C-D**). The triangulation ratio (**Fig. 2E**) and rise time (**Fig. 2F**) were significantly increased in cardiomyocytes treated with *P.a.* C-media compared to control cardiomyocytes. Together, these data show that C-media from *P.a.* infected hMDMs prolonged the AP of cardiomyocytes, as demonstrated by increased APDs and the increased percentage of beats with EAD. Furthermore, the increased triangulation ratio and time rise confirmed that prolonged AP causes arrhythmia in cardiomyocytes treated with *P.a.* C-media.

**Figure 2:**
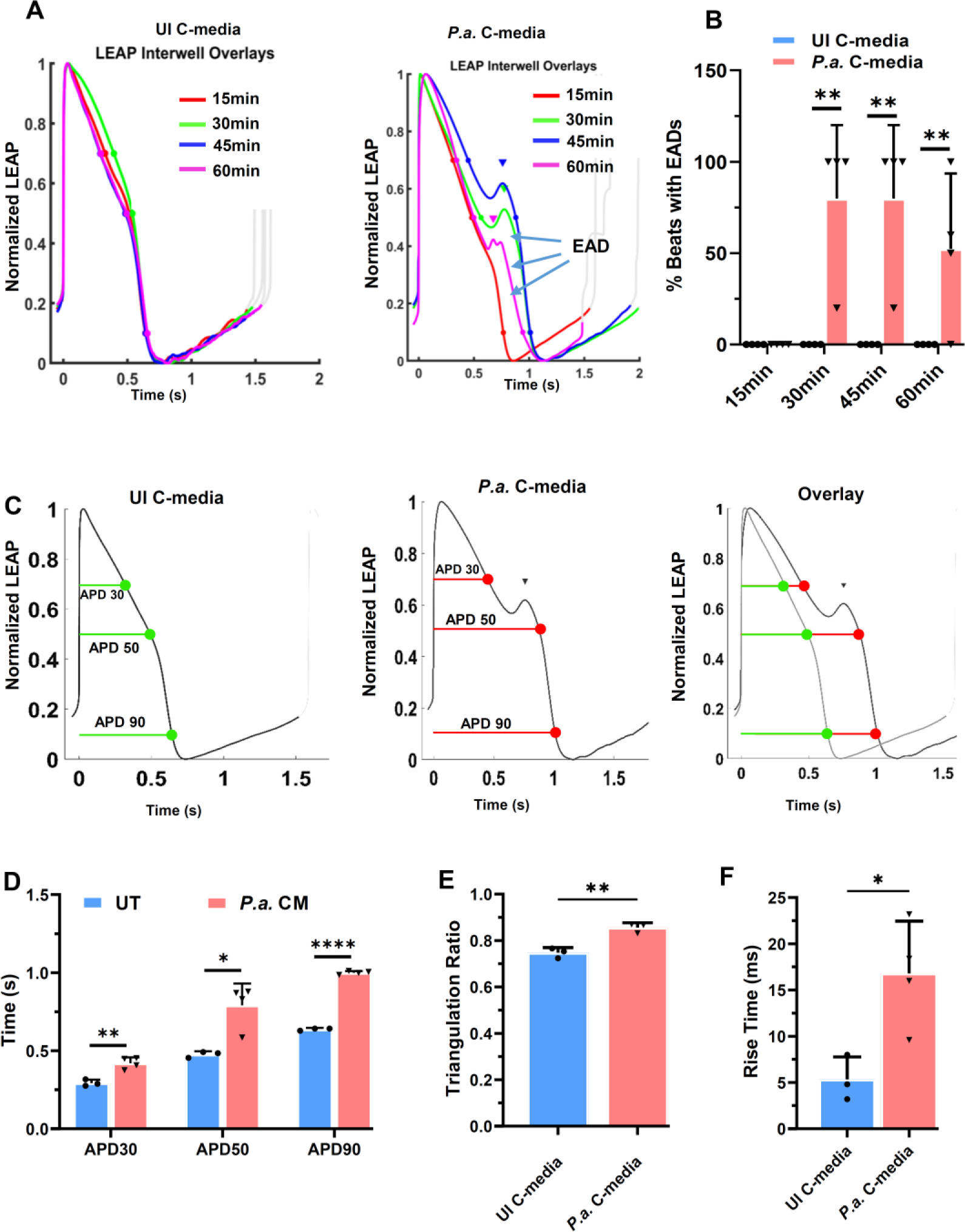
***P.a.* C-media collected from hMDMs induces arrhythmia in hiPSC-CMs.** hiPSC-CMs were cultured on the 24-well MEA plate (30x10^3^ cells/well for 7-10 days for synchronization. The hiPSC-CMs were exposed to C-media from uninfected and *P.a.* infected hMDMs and recorded the physiological parameters of cardiomyocytes on MEA at 15-, 30-,45-, and 60 min. (**A)** The action potential with induction of EAD shoulders at different time points. (**B)** The % beats with EAD features. (**C)** The representative action potential durations (APDs) at different repolarization states (APDs 30, 50, and 90). **(D)** Collective APDs at different repolarization (**E**) triangulation ratio and (**F**) rise time of depolarization of hiPSC-CMs treated with UI C-media and *P.a.* C-media. Data is representative of three independent experiments, mean ± SD: * p<0.05, ** p<0.01, **** p < 0.0001.

### *P.a.* infection enhances the release of inflammatory cytokines and extracellular vehicles (EVs) from hMDMs

Infection of hMDMs induces a robust host response resulting in enhanced production of inflammatory cytokines and release of micro vesicles into the media^38–40^. Therefore, we have sought to explore the levels of inflammatory cytokines and EVs in the C-media taken from *P.a.* infected hMDMs. First, we determined the levels of inflammatory cytokines released by hMDMs in response to *P.a.* infection. Our data indicate that *P.a.* infection enhances the release of inflammatory cytokines such as IL-1β, IL-6, and TNFα (**Fig. 3A-C**). Interestingly, we found that *P.a.* infection also induces the production of anti-inflammatory cytokine IL-10 in hMDMs compared to uninfected cells (**Fig. 3D**). Next, we quantified the EVs in the C-media using Nano Analyzer. The results showed that *P.a.* infection significantly enhances the release of EVs and showed the size and number of EVs are increased significantly by infection (**Fig. 3E-G**). Since the level of inflammatory cytokines and EVs was significantly higher in the C-media harvested from *P.a.* infected hMDMs, we next asked whether the inflammatory cytokines in the C-media causes cardiomyocyte contractile dysfunction. To evaluate this question, we incubated the C-media at 65°C for 45 min. After cooling to room temperature, we incubated C-media with hiPSC-CMs and monitored their electrophysiology. We used non-heat-inactivated C-media as a control. Surprisingly, our MEA data showed that the addition of heat-inactivated C-media from *P.a.* infected hMDMs to hiPSC-CMs caused quick and rapid contractile dysfunction compared to non-heat-inactivated C-media exposure (**S**.**Fig. 2A**). Also, we determined the beat period **(S.Fig. 2C**), beat rate, and FPD (**S**.**Fig. 2D-E**). Our data showed that in comparison to non-heat-inactivated C-media, heat-inactivated C-media rapidly decreased the beat rate (**S**.**Fig. 2D**) and prolonged the depolarization and repolarization duration as shown by increased FPD in this group (**S.Fig. 2E**). Thus, the data indicates that at elevated temperatures, the EVs could release the content (bacterial PAMPs) into the medium and cause rapid cardiomyocyte dysfunction.

**Figure 3:**
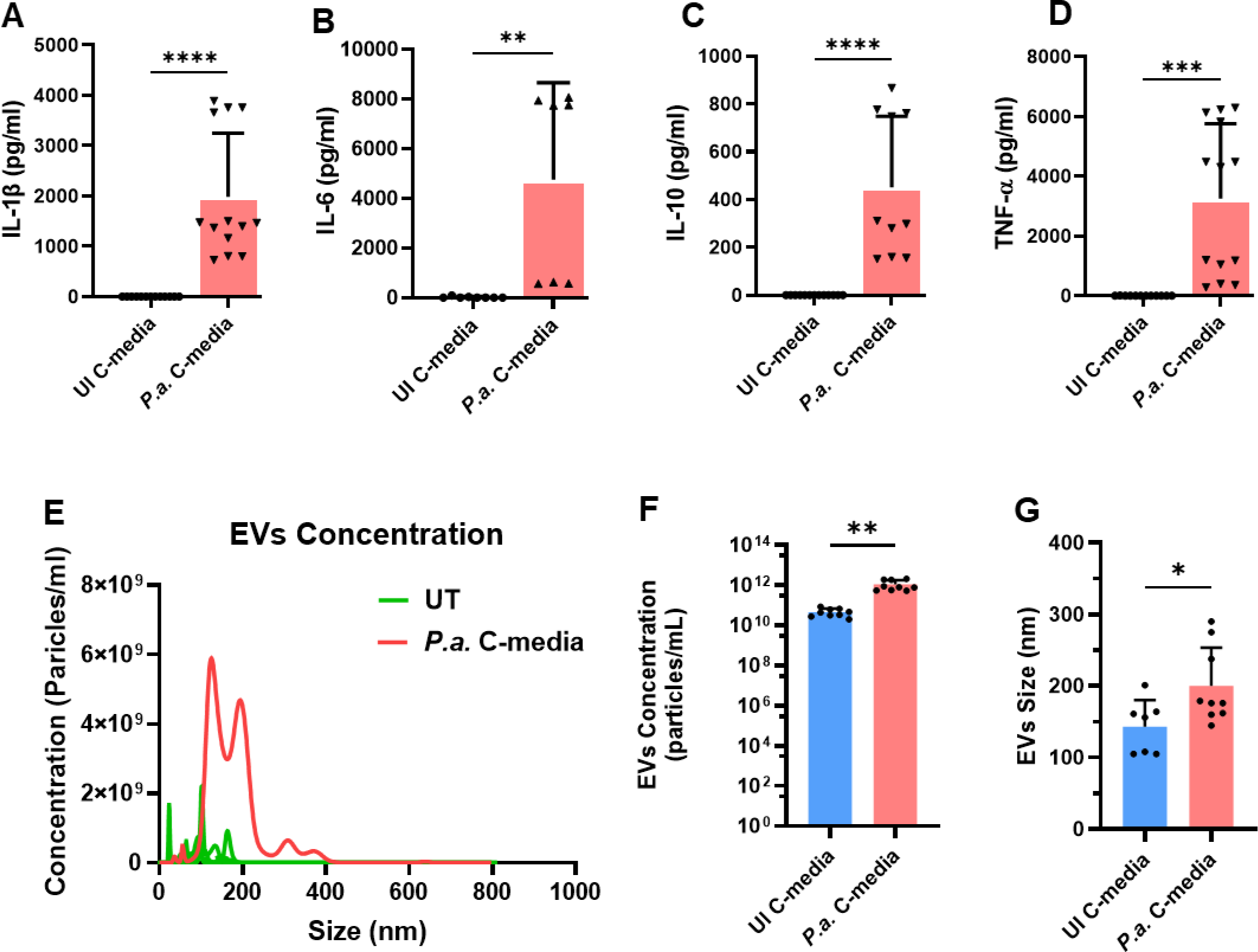
P*.a.* infection enhances the release of cytokines and exosomes. hMDMs were infected with *P.a.* (1 MOI) or left uninfected for 22h. The cell free culture supernatants were evaluated by ELISA to determine the cytokine production. Graphs shown are (**A**) IL-1β, (**B)** IL-6, (**C)** IL-10, and (**D)** TNF-α levels in C-media. Exosomes isolated from UI-C-media and *P.a.* C-media were quantified using NanoFCM. The plot shown in (**E**) is the distribution of EV size versus their concentration. (**F)** EVs concentration, and (**G)** EVs size. Accumulative data from three independent experiments, mean ± SD: ** p <0.01, *** p< 0.001, **** p < 0.0001.

### Live *P.a.* is required to cause cardiac dysfunction in hiPSC-CMs

Next, we asked whether live bacterial infection is necessary to induce the production of inflammatory cytokines and EVs in hMDMs. We infected hMDMs with live *P.a.* and heat-killed *P.a.* After 22-hr of incubation, C-media was harvested and exposed to hiPSC-CMs, and contractile function was monitored using the MEA system. These data revealed that live bacterial infection of hMDMs is necessary to induce cardiomyocyte contractile dysfunction (**Fig. 4A**). Also, we observed that C-media harvested from live *P.a.* infected hMDMs caused an irregular beat period, decreased beat rate, and FPD (**Fig. 4B- D**). However, the C-media harvested from heat-killed *P.a.* infected hMDMs did not affect the cardiomyocyte contractile function. Next, we determined the levels of cytokines in the C-media harvested from live and heat-killed *P.a.* infected hMDMs. The ELISA data revealed that incubation of hMDMs with heat-killed *P.a.* releases elevated levels of IL-1β, IL-6, TNFα, and IL-10 compared to live *P.a*. infected hMDMs (**Fig. 4E-H**), due to the increased cell death in live P.a. infection. Thus, two conclusions can be drawn from these studies. First, the cardiomyocyte contractile dysfunction requires active *P.a.* infection of hMDMs and second, inflammatory cytokines in the C-media harvested from infected hMDMs are not sufficient to induce cardiomyocyte contractile dysfunction.

**Figure 4:**
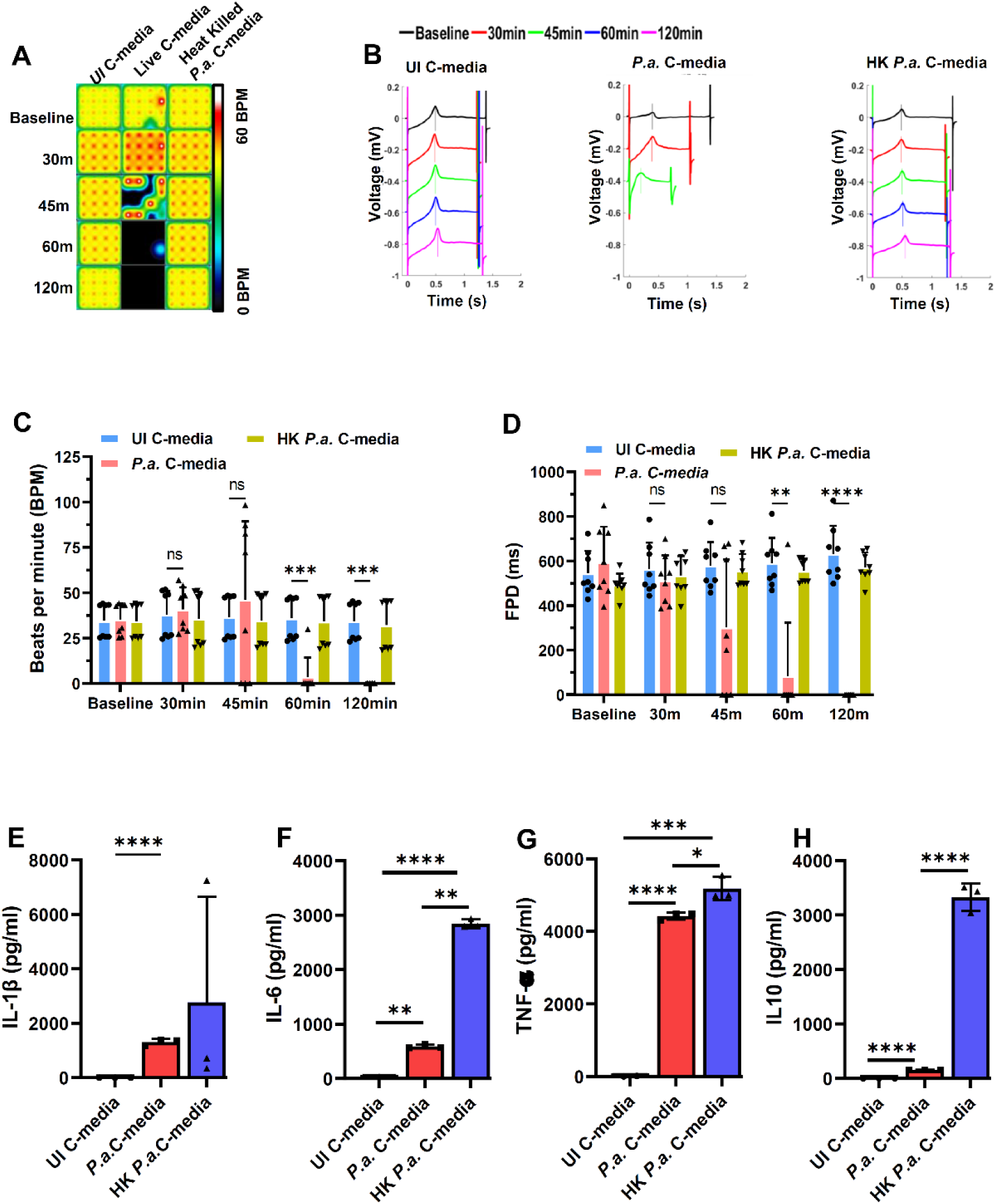
Live bacteria are required to cause cardiomyocyte contractile dysfunction. hiPSC-CMs were plated in 24 well MEA plates, and the cells exposed to the mixture of cardiomyocyte culture medium and C-media (1:1 ratio) harvested from uninfected (UI C-media), *P.a.* infected (*P.a.* C-media) or heat-killed *P.a.* infected (HK *P.a*. C-media) hMDMs. The cardiomyocyte contractility and electrical activity was recorded using AxIS Navigator on the MEA system at 5% CO2 and at 37°C for indicated time points. Data analysis was performed using the cardiac analysis tool. (**A**) Electrical activity map showing the changes in beat rate of cardiomyocytes treated with UI C-media and *P.a.* C- media. The activity map shown is a representative well from quadruplicate samples for each treatment and three repeats (N=3). (**B**) Representative traces recorded with the MEA system showing the beat period, T-wave, and FPD in UI-C-media, *P.a.* C-media, and HK *P.a.* C-media-treated hiPSC-CMs. The graph shown in (**C**) is beats per minute, and (**D**) filed potential duration (FPD) at baseline, 30-, 45-, 60- and 120 min post-treatment of hiPSC-CMs with UI C-media, *P.a.* C-media, and HK *P.a.* C-media. Graphs shown in are (**E**) IL-1β, (**F)** IL-6, (**G)** TNF-α, and (**H)** IL-10 levels in C-media harvested from uninfected, live *P.a.* and heat-killed *P.a.* infected hMDMs. Data shown in Figures C and D are accumulative data from 3 independent experiments (mean ± SD; ** p <0.01, *** p < 0.001, **** p < 0.0001). Data shown in figure E-H is representative data from two independent experiments, mean ± SD: * p<0.05, ** p <0.01, *** p < 0.001, **** p < 0.0001.

We predict that danger-associated molecular pattern (DAMPs) released from infected host cells and pathogen-associated molecular patterns (PAMPs) released from live bacteria may play critical roles in causing cardiomyocyte damage. To evaluate whether PAMPs or DAMPs are responsible for cardiomyocyte contractile dysfunction, we exposed the hiPSC-CMs to C-media harvested from *P.a.* infected hMDMs and *P.a.* cultured in RPMI media supplemented with 10% serum and monitored the cardiomyocyte electrical activity. Our MEA data revealed that C-media from *P.a.* cultured in RPMI media also caused hiPSC-CM contractile dysfunction by altering their electrical activity and decreasing their beat rate (**S.Fig. 3A-B**) to the same extent as C-media harvested from *P.a.* infected hMDMs. These data suggest that PAMPs from bacteria are primarily responsible for causing cardiomyocyte contractile dysfunction.

### OMVs released from *P.a.* induce cardiomyocyte contractile dysfunction

It has been known that *P.a.* secretes bacterial products, including toxins and OMVs^41,42^. These OMVs are packed with bacterial toxins and antigens^42^. To test whether *P.a.* OMVs cause cardiomyocyte contractile dysfunction, we grew *P.a.* in LB culture media for 22h and isolated OMVs from the culture media using published methods^42^. We exposed hiPSC-CMs to OMVs (1:3.8x10^6^ ratio hiPSC-CMs: OMVs) or left them untreated and monitored cardiomyocyte contractile function. The MEA data showed that exposure of hiPSC-CMs to OMVs significantly increased their beat period (**Fig. 5A**) and decreased their beat rate (**Fig. 5B**). Next, we determined the FPD, which represents the duration between depolarization and repolarization of the cardiomyocytes. Our data showed a significant decrease in the FPD of cardiomyocytes treated with OMVs compared to untreated cardiomyocytes (**Fig 5C**). Furthermore, we observed that treatment of hiPSC-CMs with OMVs causes irregular beat periods over a three hundred-second duration (**Fig. 5D vs. 5F**). We further analyzed the individual beats present during this period and found that as compared to the untreated group (**Fig. 5E**), in cardiomyocytes treated with OMVs, their beats were at irregular intervals with missing depolarization spikes **(Fig. 5G**). Thus, the data show that like *P.a.* C-media treatment (**Fig. 1F-I**), treatment of hiPSC-CMs with OMVs also causes arrhythmia.

**Figure 5:**
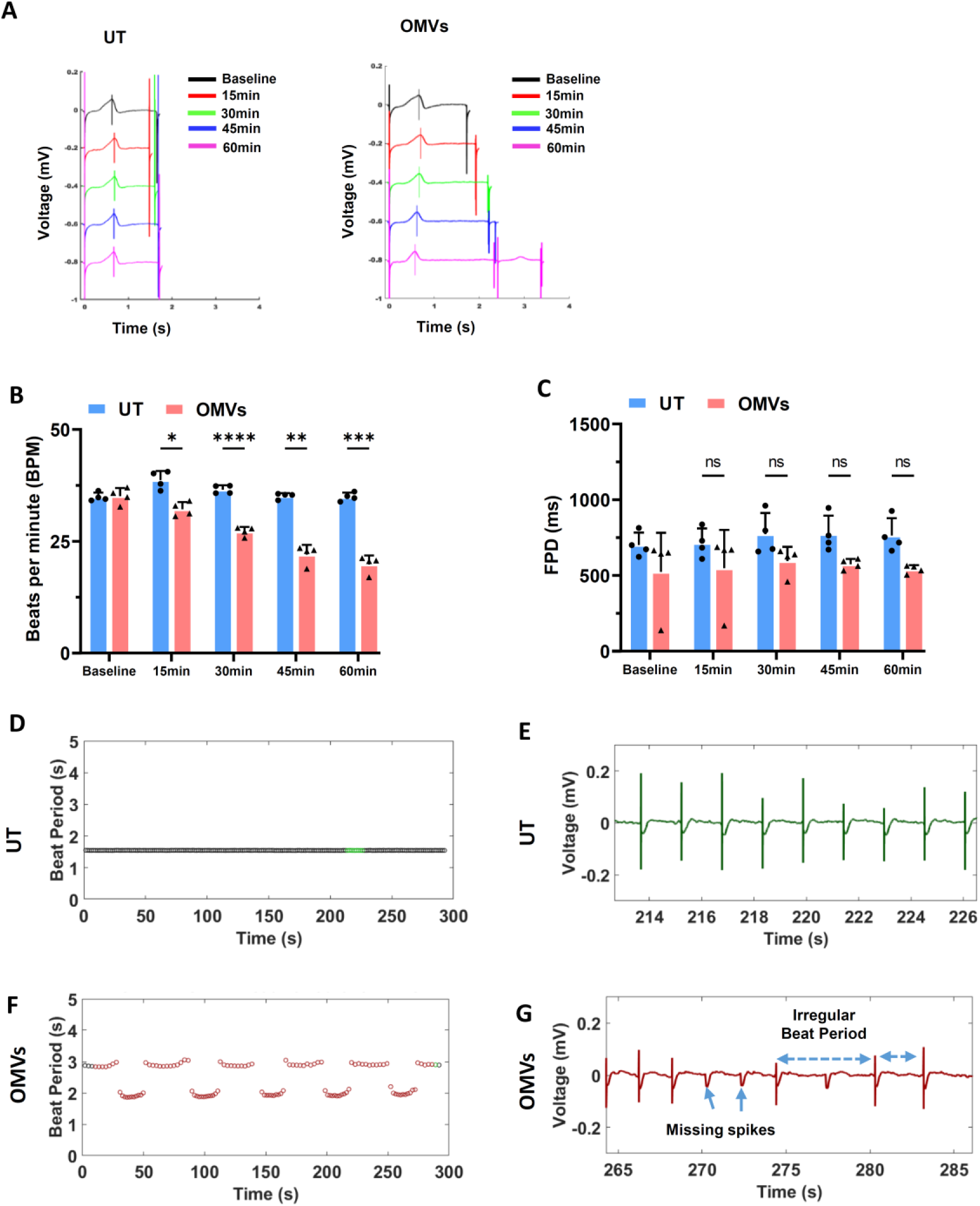
P*.a.* OMVs are potent inducer of cardiomyocyte contractile dysfunction. hiPSC-CMs were cultured on the MEA plate and exposed with OMVs or left untreated (UT), and the cardiomyocyte contractility and electrical activity were recorded using AxIS Navigator on the MEA system at 5% CO2 and at 37°C for indicated time points. (**A**) Representative traces recorded with the MEA system showing the beat period, T- wave, and FPD in UT and OMV-treated hiPSC-CMs. The graphs shown in (**B**) are beats per minute and (**C**) FPD at baseline, 15-, 30-, 45-, and 60 min post-treatment of hiPSC-CMs with OMVs. (**D-E**) Representative beat period of untreated (**F-G**) OMV treated hiPSC- CMs over time. Data shown in Figures B and C are representative of 4 independent experiments, mean ± SD: * p<0.05, ** p <0.01, *** p < 0.001, **** p < 0.0001.

### *P.a.* OMVs and conditioned medium from *P.a.* infected hMDMs inhibit calcium transient cardiomyocytes and causes arrhythmia

Intracellular Ca^2+^ cycling is crucial for regulating cardiac rhythms, and any disturbances in Ca^2+^ handling can cause arrhythmias^43,44^. Since we observed that exposure of hiPSC-CMs with OMVs caused severe cardiomyocyte contractile dysfunction, we investigated its impact on cellular Ca^2+^. hiPSC-CMs were stained with Fluo-3 AM to evaluate the intracellular Ca^2+^ dynamics in the presence of OMVs harvested from *P.a.* or left untreated. OMV treatment significantly delayed the intracellular Ca^2+^ transients compared to the control in hiPSC-CMs, which are observed in Ca^2+^ transient traces (**Fig. 6A-C)**. The peak amplitude of cytosolic Ca^2+^ was significantly elevated in OMV-treated hiPSC-CM compared to the control **(Fig. 6D)**. As shown in **Fig. 6E**, an irregular and longer Ca^2+^ cycling duration was observed under OMV treatment compared to the control. Also, the time-to-peak was decreased and decay time was significantly increased in the OMV treatment group compared to the control (**Fig. 6F and G**). These results indicate that OMVs affect oscillations of intracellular Ca^2+^ in cardiomyocytes, leading to arrhythmogenic phenotype.

**Figure 6:**
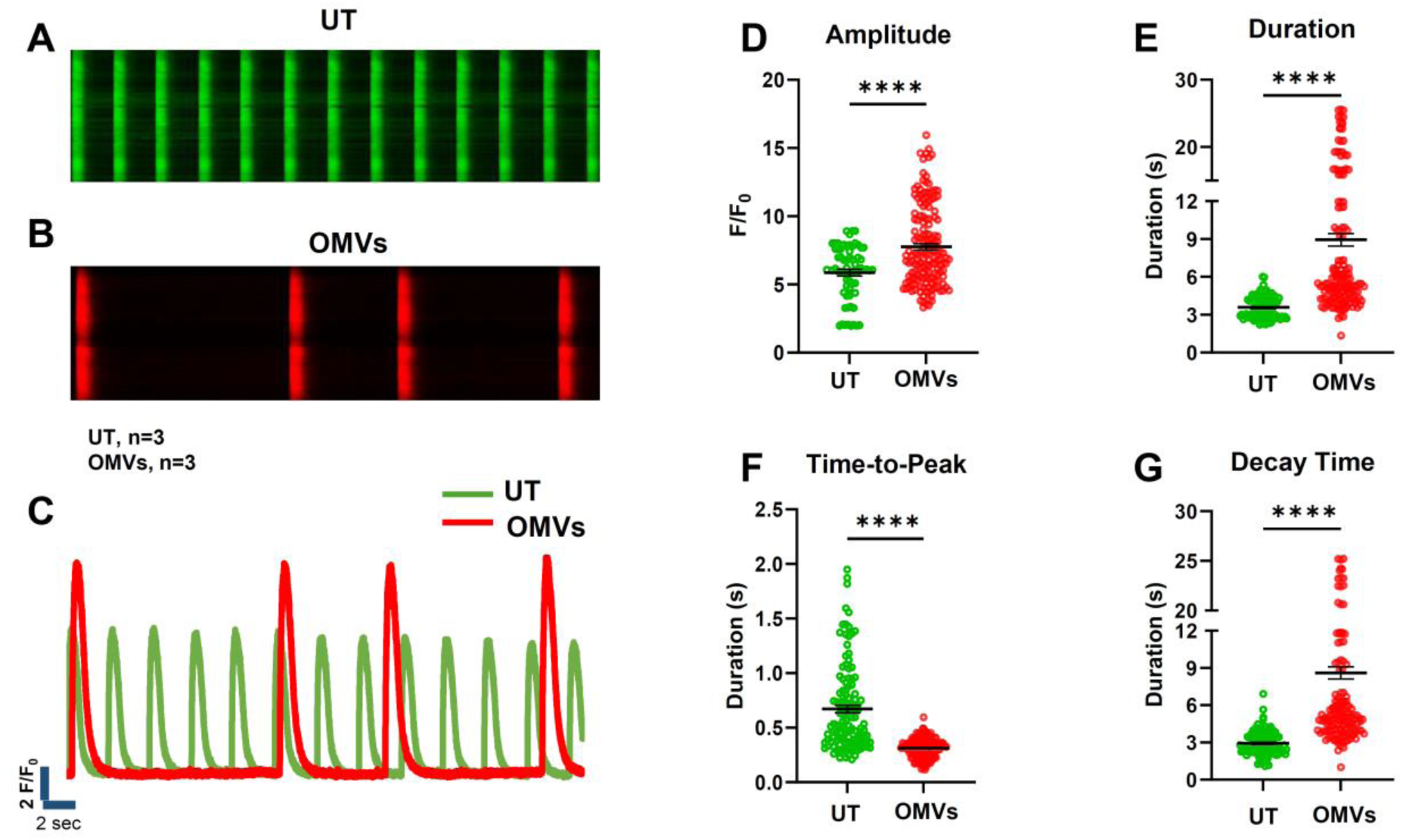
C-media and bacterial OMVs dysregulate the calcium channel function in cardiomyocytes. Synchronized hiPSC-CMs were treated with UI-C-media and or *P.a.* C-media, and the Ca²⁺ transient was visualized using Fluo-3AM. Representative images of Ca^2+^ transient wave of cardiomyocytes exposed to UI-C-media (**A**) and *P.a.* C-media (**B**). (**C**) Representative overlay of Ca^2+^ traces collected from cardiomyocytes incubated with UI-C-media and *P.a.* C-media. Ca^2+^ transient parameters (**D**) peak amplitude, (**E**) peak duration, (**F**) time- to Peak duration, and (**G**) decay time were measured in UI C-media and *P.a.* C-media treated cardiomyocytes. (**H-I)** Representative traces showing Ca^2+^ transient wave of cardiomyocytes left untreated or exposed to *P.a.* OMVs. (**J**) Representative overlay of Ca^2+^ traces collected from cardiomyocytes incubated with OMVs. Ca^2+^ transient parameters (**K**) peak amplitude, (**L**) peak duration, (**M**) time- to Peak duration, and (**N**) decay time were measured in UI and OMV-treated cardiomyocytes. Graphs shown are accumulative data acquired from 10-15 cells of three independent experiments; a Two-tailed Student’s t-test was performed for comparison with control, **** p˂0.0001.

Since, we found that the C-media induces early after-depolarizations (EAD) and severe arrhythmia (**Fig. 1 and 2**), we investigated its impact on cellular Ca^2+^. hiPSC-CMs were stained with Fluo-3 AM to evaluate the intracellular Ca^2+^ dynamics in the presence of C-media harvested from *P.a.* infected hMDMs compared with the cardiomyocyte medium (**S.Fig. 4A-C**). Spontaneously beating cardiomyocytes showed a significant increase in peak amplitude within 15 mins of treatment with C-media compared to untreated cardiomyocytes (**S.Fig. 4D)**. Similarly, hiPSC-CMs treated with C-media showed irregular Ca^2+^ duration intervals as compared to the untreated hiPSC-CMs, which suggests that C-media disrupts the synchronous beating of cardiomyocytes (**S.Fig. 4E).**

Likewise, the Ca^2+^ transient parameters, time-to-peak was decreased and decay time was significantly increased by C-media treatment compared to untreated hiPSC-CMs (**S.Fig. 4F-G**). These findings suggest that the inflammatory mediators or other components in the C-media negatively affect Ca^2+^ dynamics in hiPSC-CMs, potentially leading to impaired contractile function and arrhythmias.

### EVs released from *P.a.* infected hMDMs are packed with bacterial components

Since, our data demonstrated that incubation of C-media harvested from P.a. infected hMDMs causes cardiomyocyte contractile dysfunction (**Fig.1 and 2).** Also, the C-media from infected hMDMs contains enormous number of EVs (**Fig. 3 E-G**). Thus, we sought to determine the contents of EVs. We purified the EVs according to the published protocol^15,45^. EVs were lysed with TN-1 lysis buffer and the lysates digested with trypsin and analyzed by LC-MS/MS Orbitrap mass spectrometry. The peptide sequences were analyzed using the MASCOT database, and the identified proteins were listed in **Table 1**. This procedure identified 295 bacterial proteins in the EVs isolated from *P.a.* infected C-media. This data suggests that EVs carry bacterial products including immunogenic proteins, toxins, and cell wall components, which are responsible for cardiomyocyte contractile dysfunction.

**Table 1:**
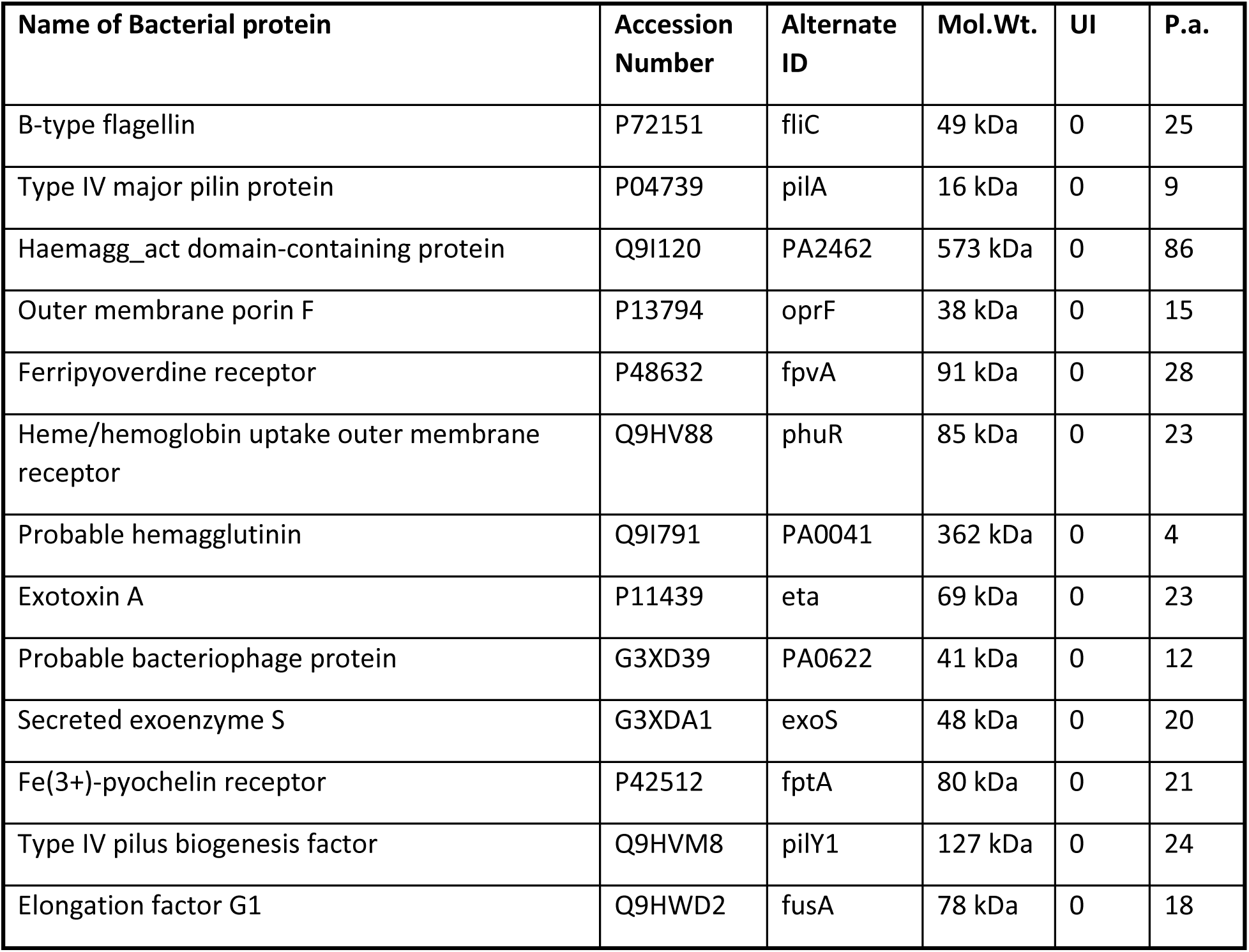
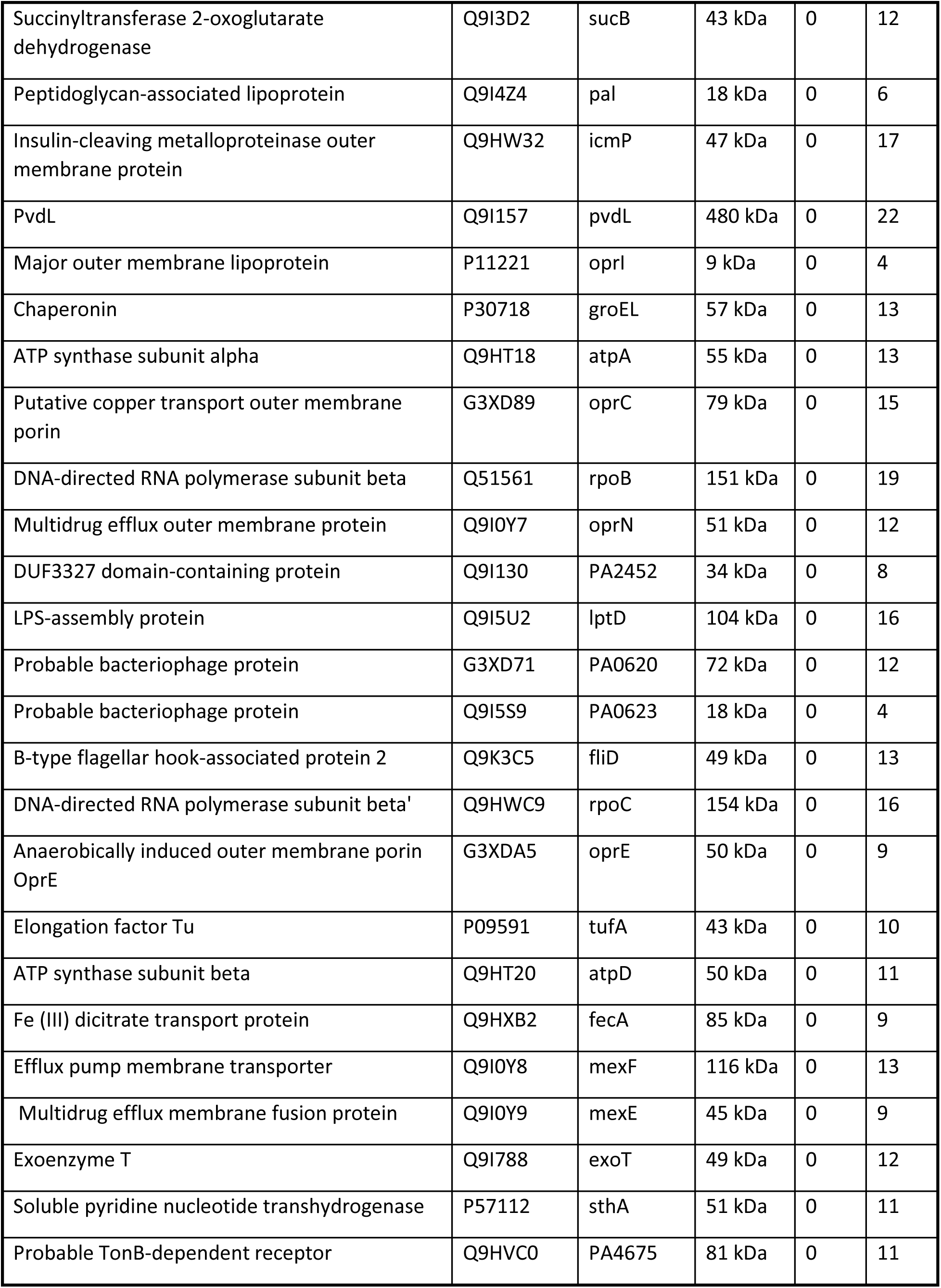
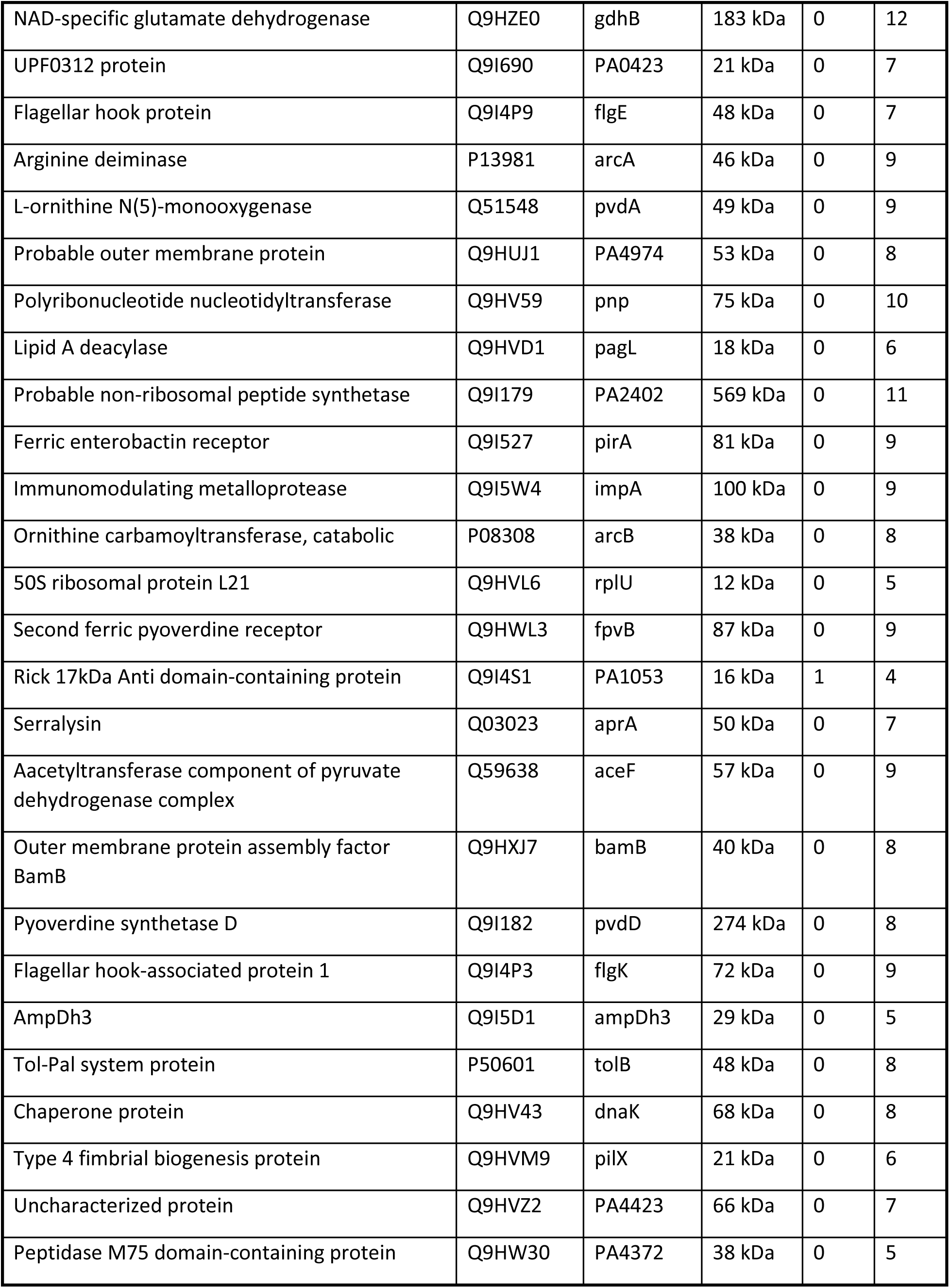
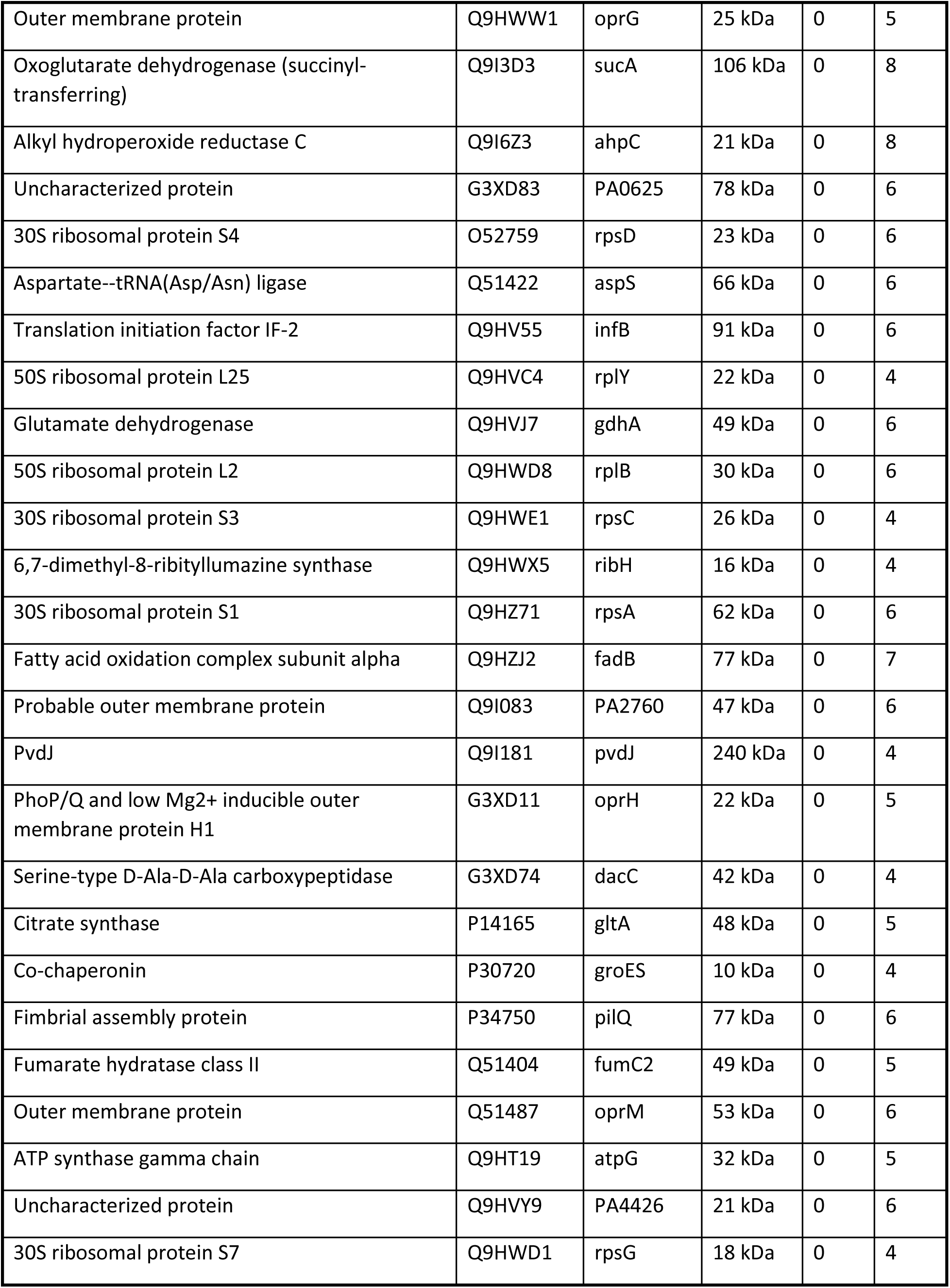
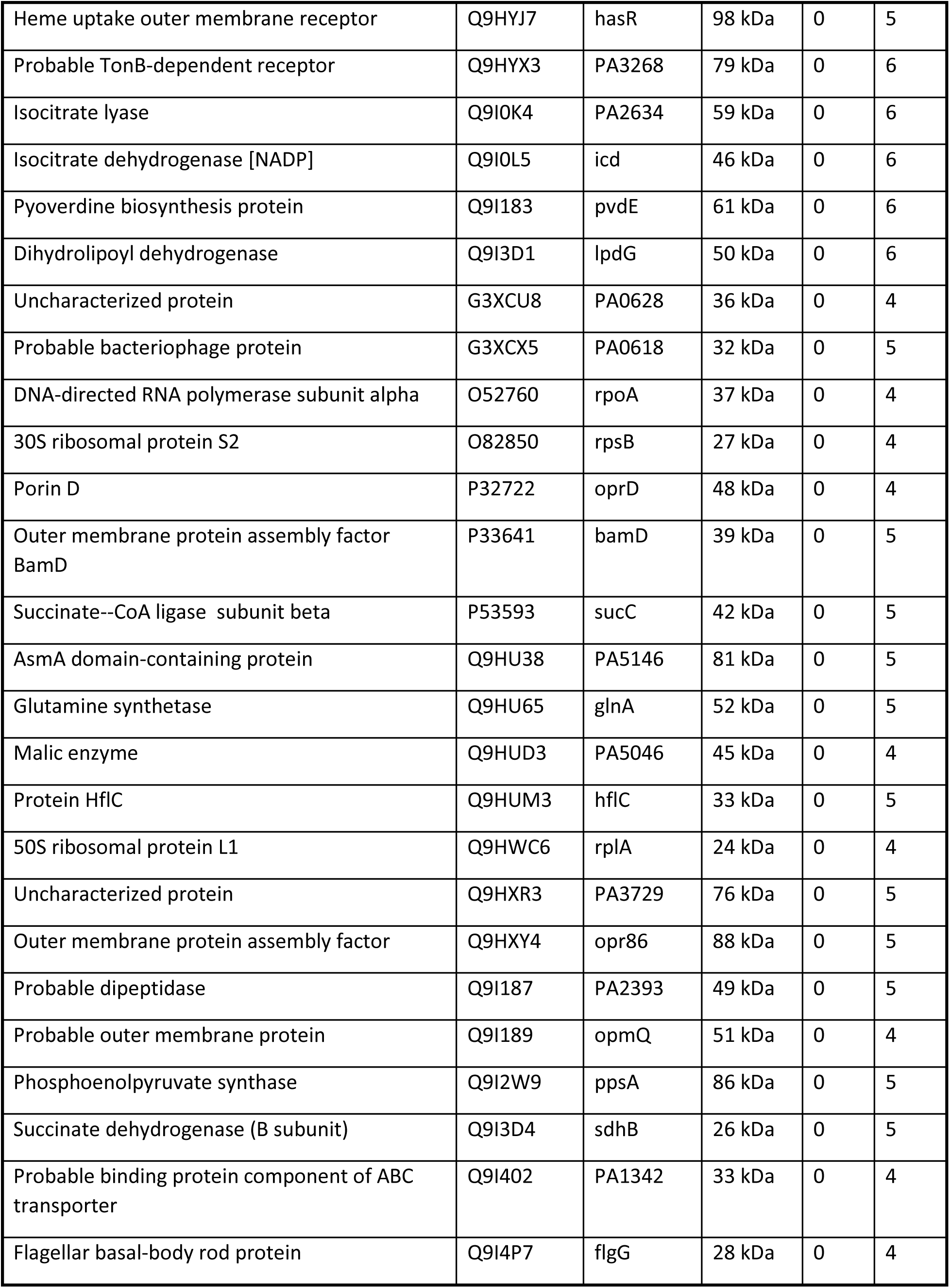
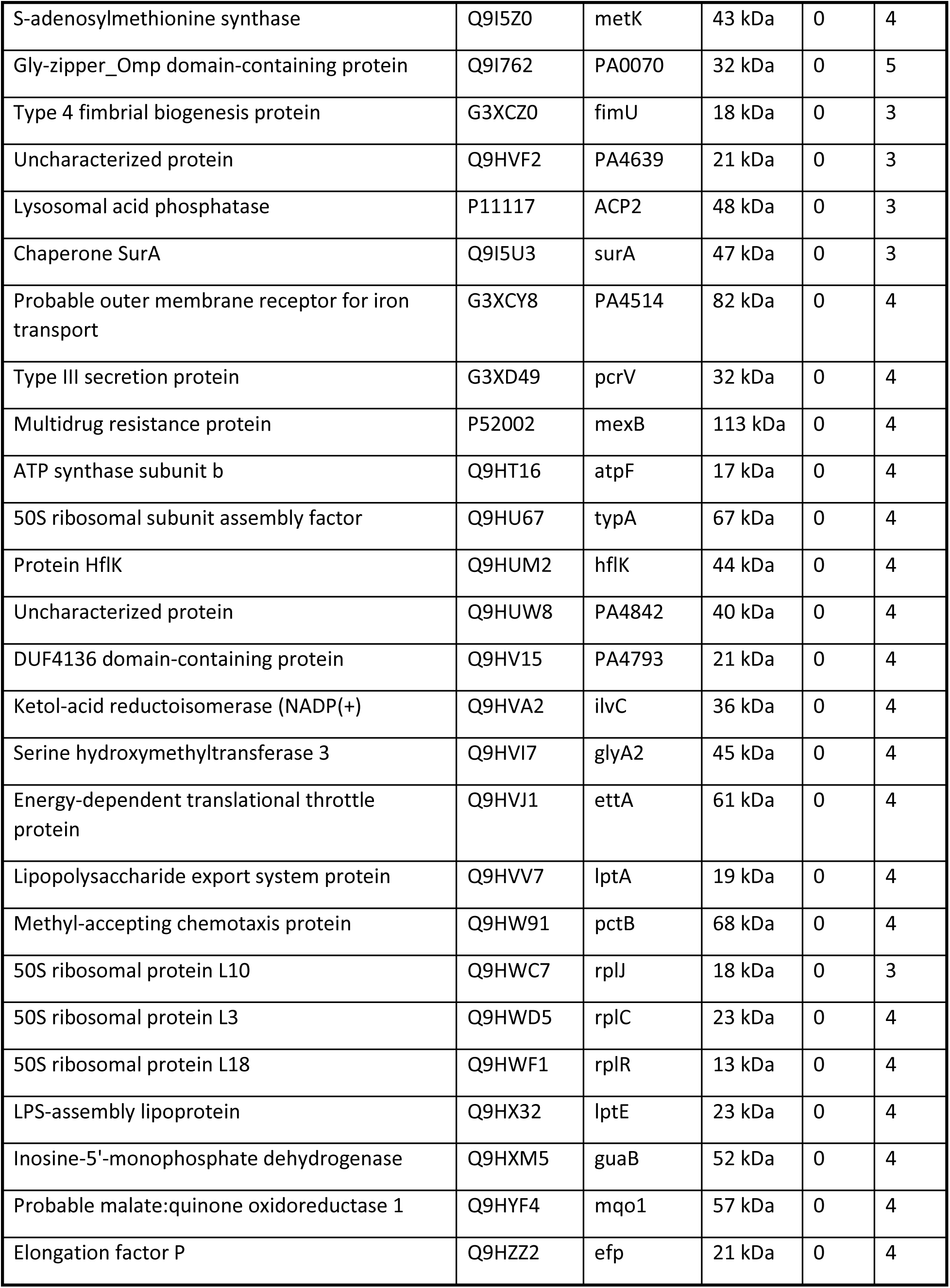
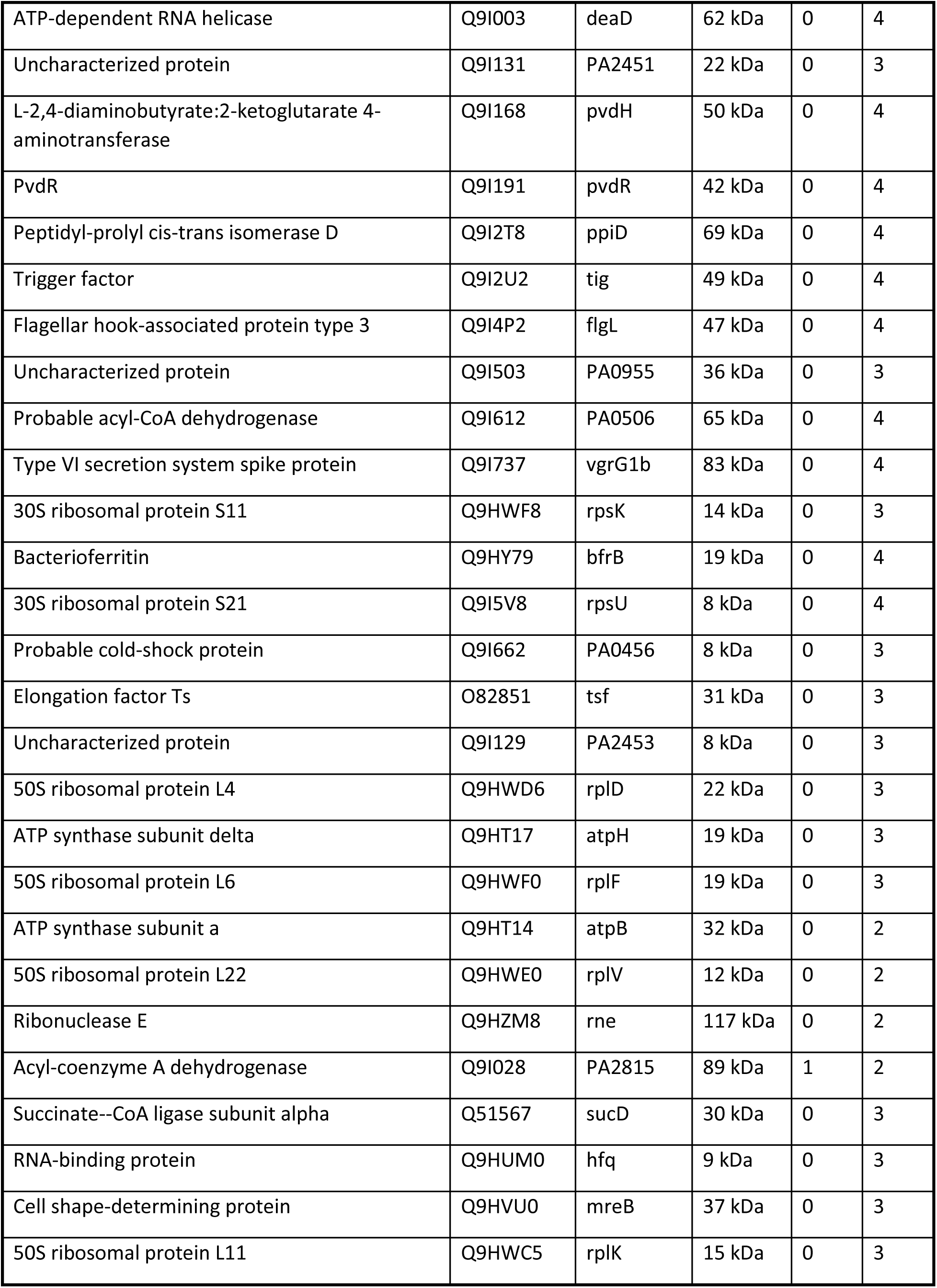
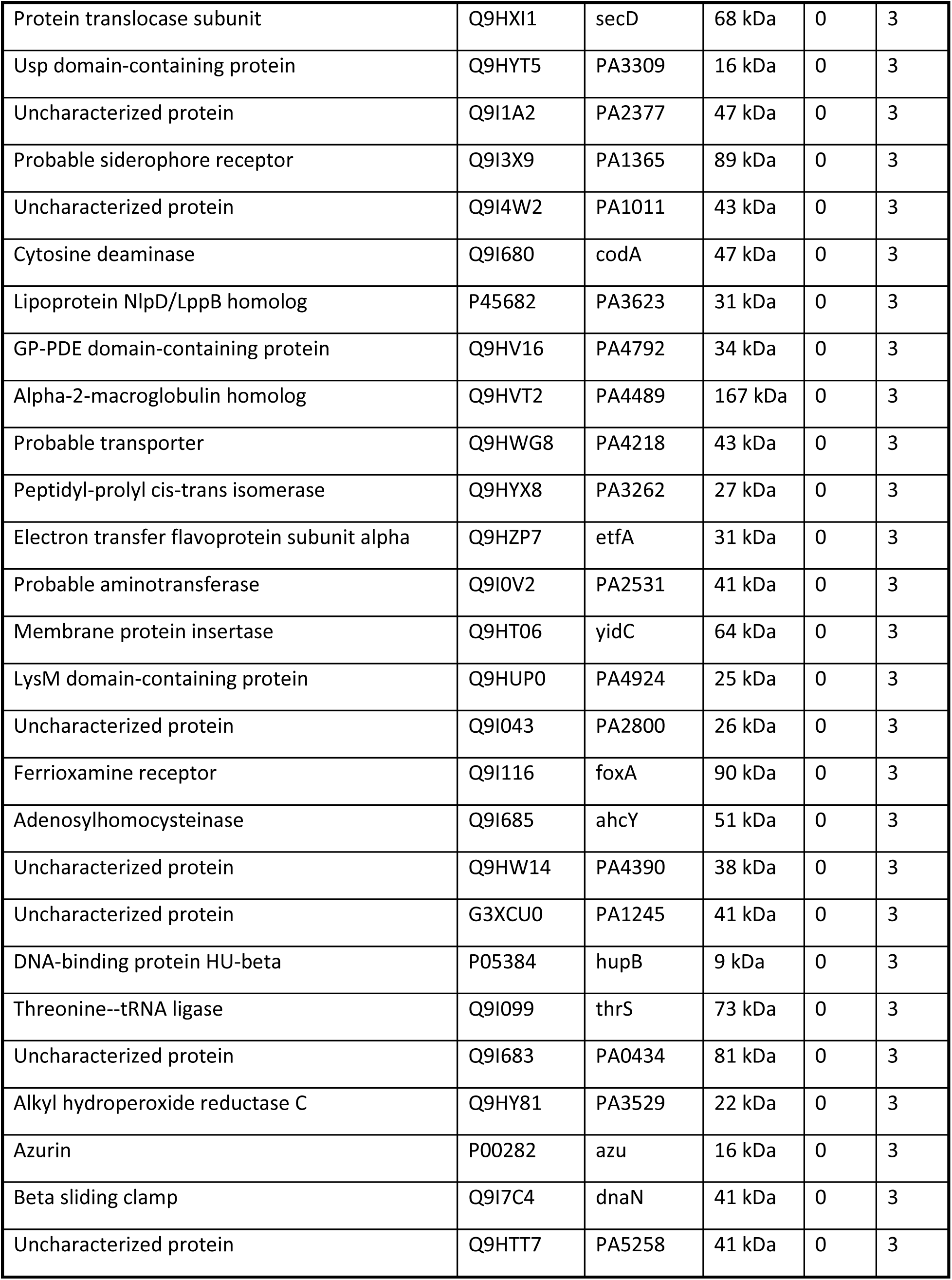
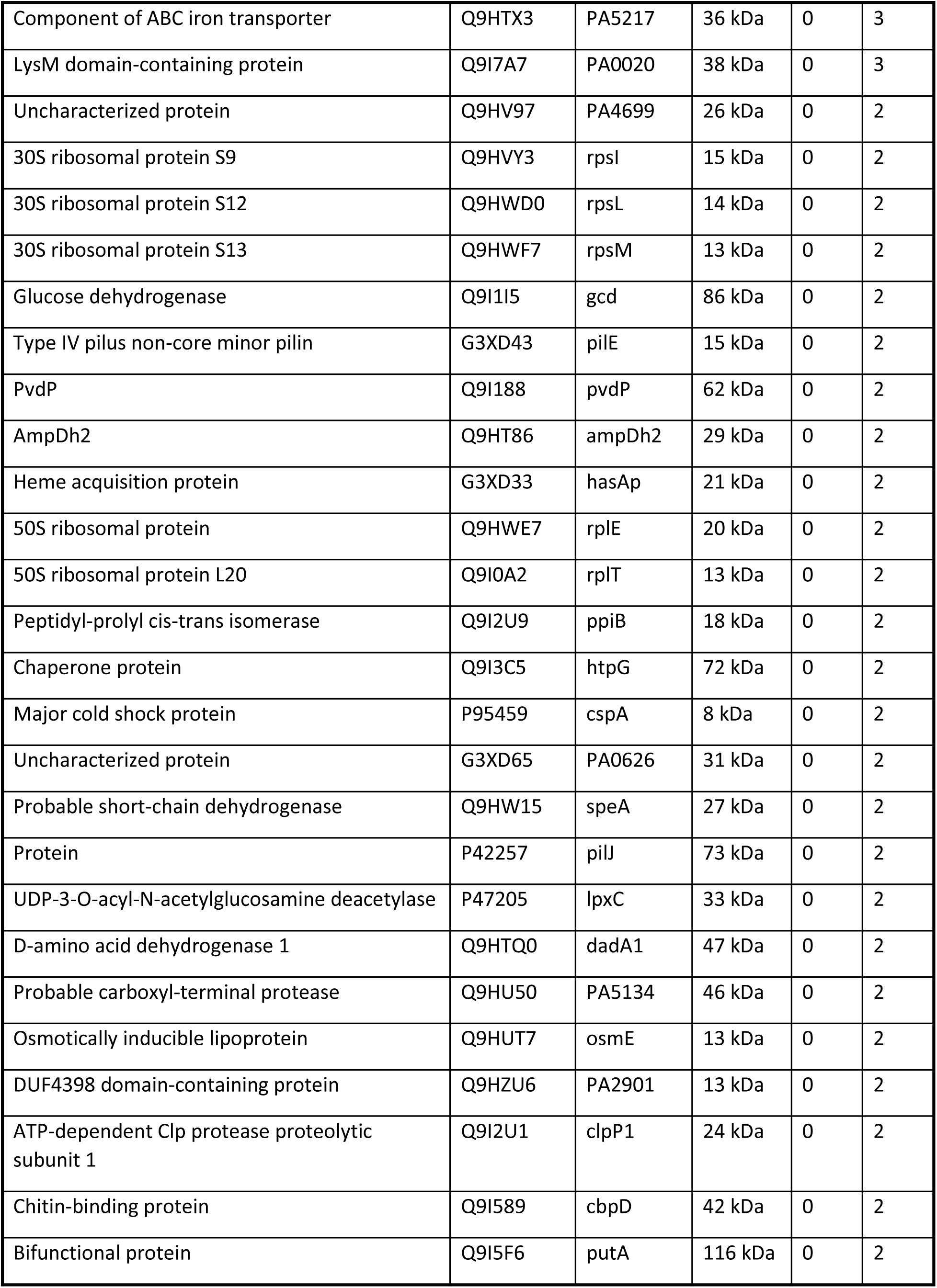
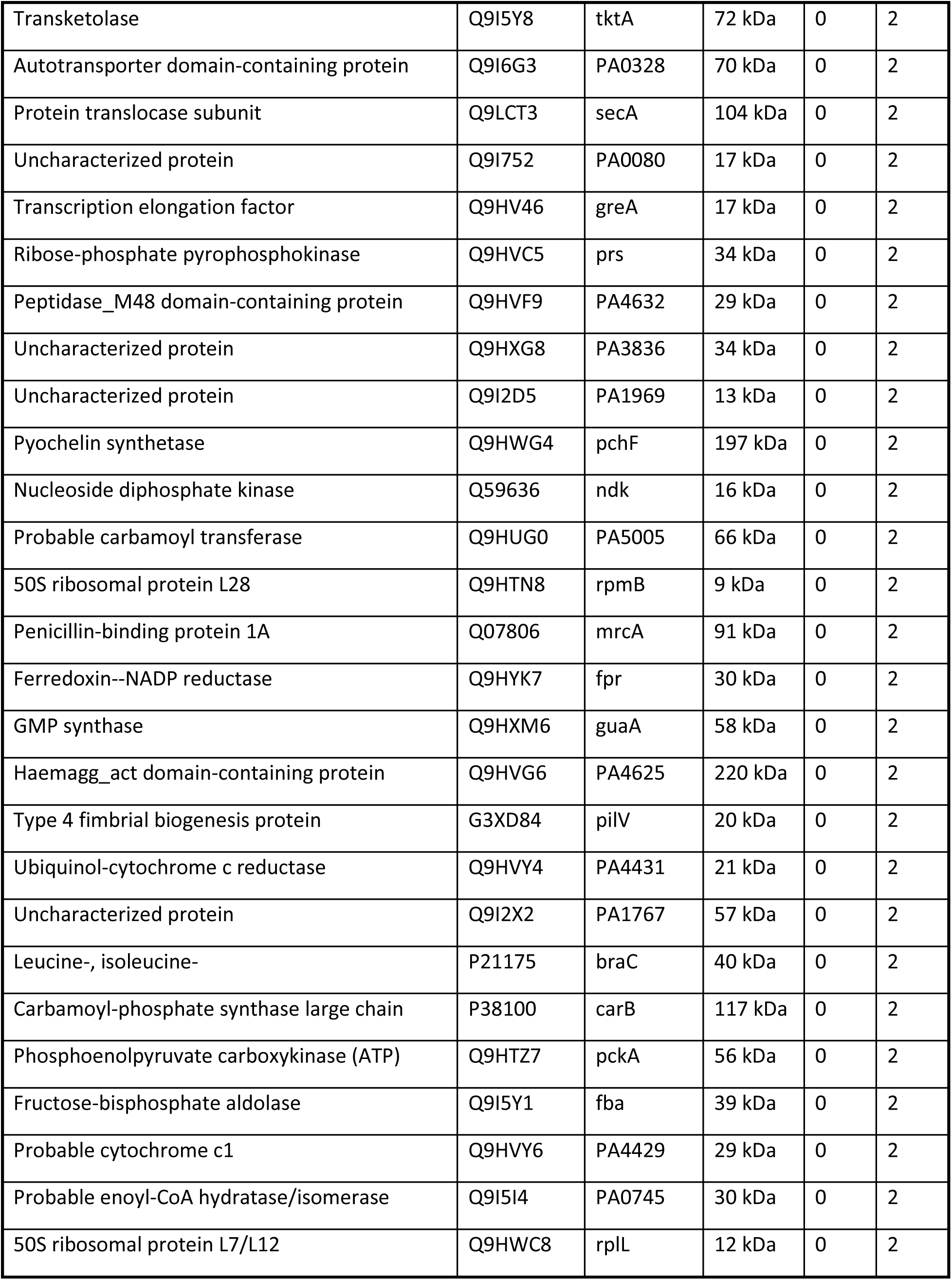
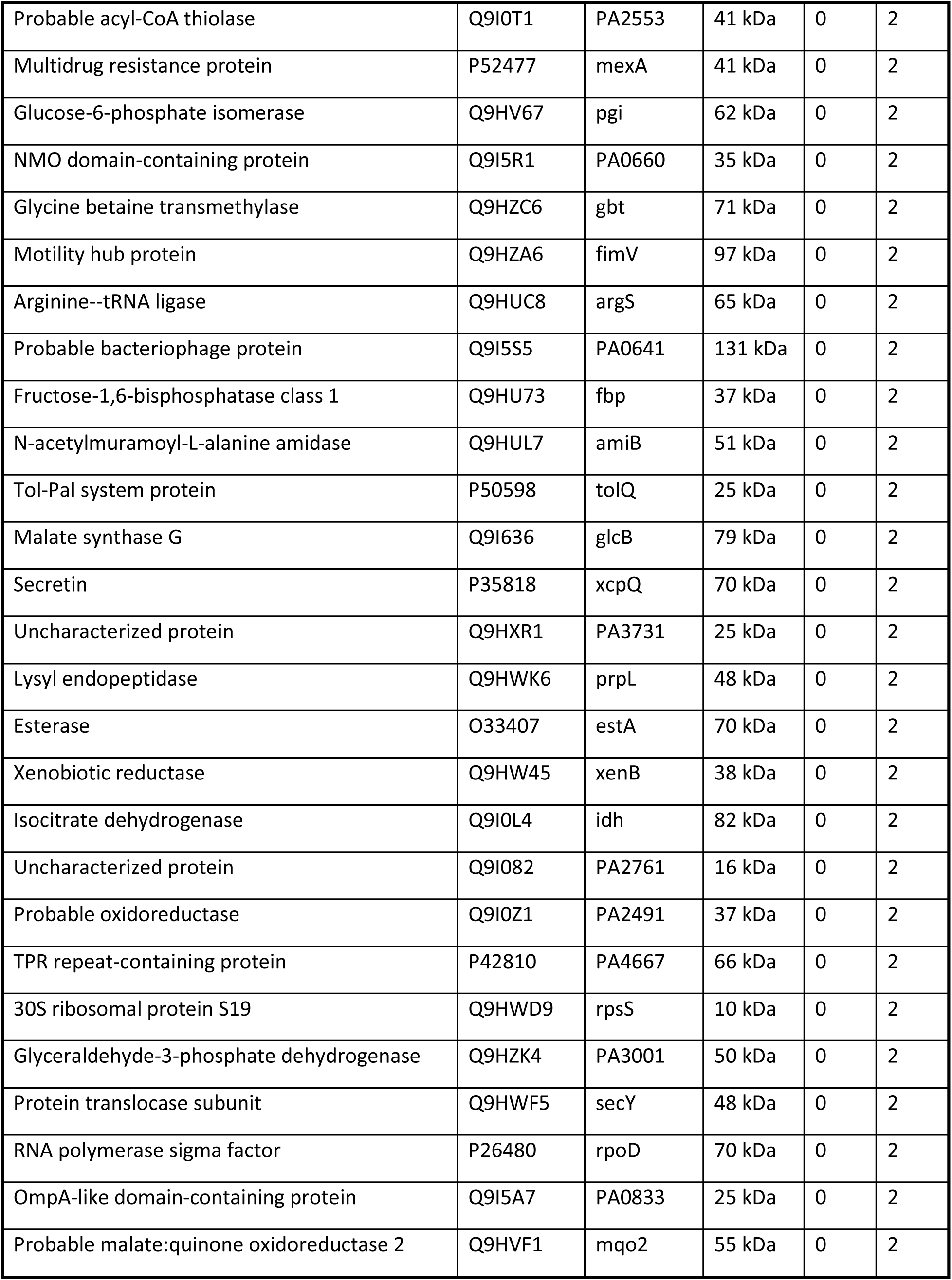
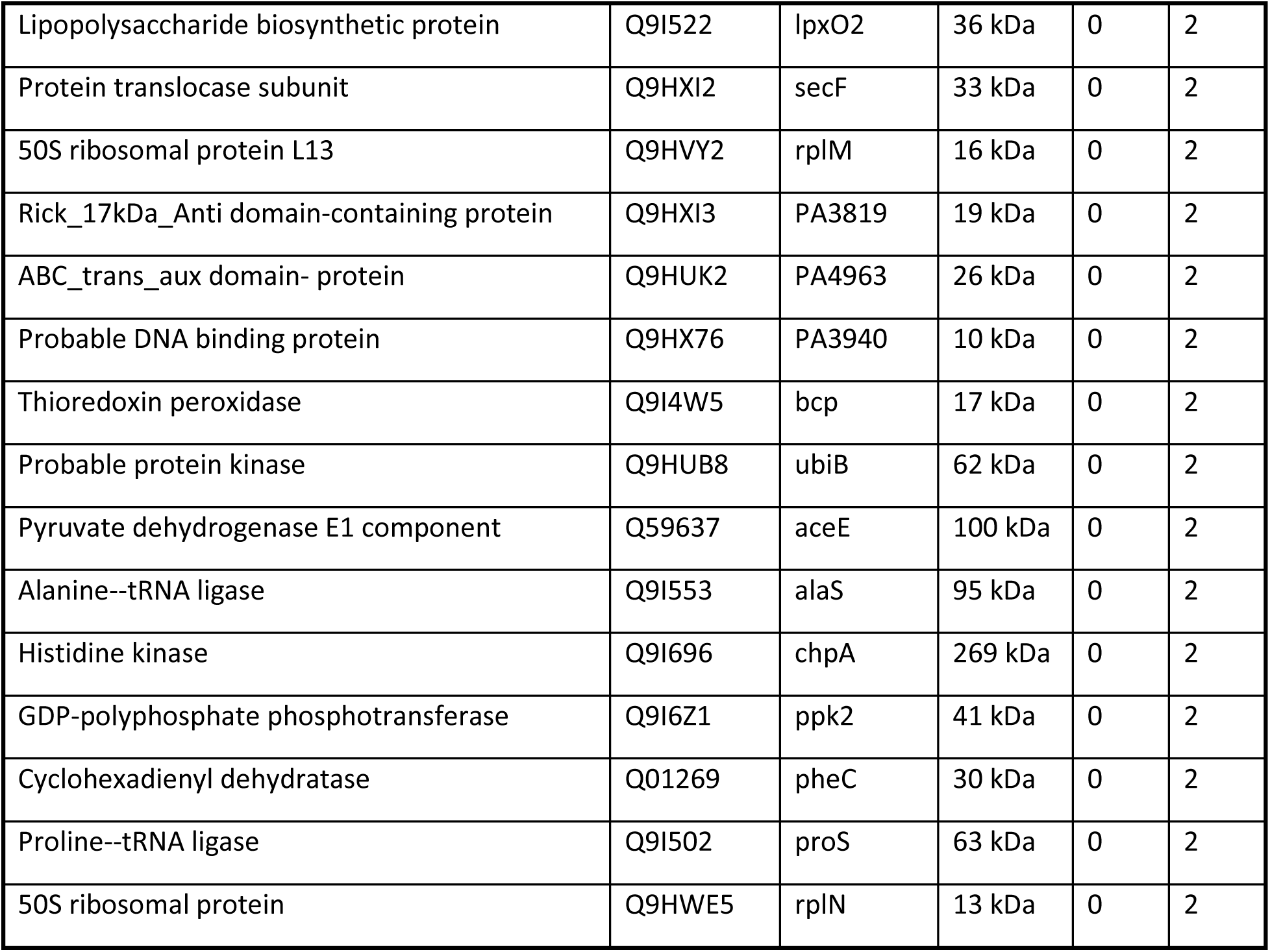
Proteins identified from exosomes harvested from uninfected (UI) and *P.a.* infected (PAO1) hMDMs by LC-MS/MS. Overnight cultured hMDMs were infected with 1 MOI of *P.a.* for 22 hours, the cell free culture supernatants were used to harvest the EVs using above-described protocol (methods section). Equal amount of EV proteins was subjected to LC-MS/MS and data was analyzed using Scaffold (^TM^) version 5.2.0. analyzed. The unique bacterial proteins found in the EVs were listed based on the presence of unique peptide score.

### EVs released from *P.a.* infected hMDMs are packed with bacterial OMVs

Our data reveal that C-media harvested from *P.a.* infected hMDMs cause cardiomyocyte contractile dysfunction and this impact cannot be explained by cytokine production (**Fig. 1, S.Fig. 2**, **Fig. 4 E-H**). Furthermore, *P.a.* C-media contains EVs (exosomes from hMDMs and OMVs from bacteria) and our LC-MS/MS data reveals that EVs contains bacterial products. However, whether the contents of the EVs come from OMVs released from *P.a.* or released bacterial products that are loaded within exosomes is not known, since the EV isolation from C-media of *P.a.* infected hMDMs, does not distinguish bacterial OMVs and host-derived exosomes. To separate the host-derived exosomes and bacterial OMVs, we incubated the purified EVs with CD9 antibody-coated magnetic beads (which specifically bind to host-derived exosomes) and separated the exosomes and bacterial OMVs (schematic shown in **Fig. 7A**). The purified exosomes and OMVs were confirmed with CD9 antibody using western blot, as shown in figure **7B**. To identify the contents of exosomes and OMVs, we subjected the CD9^+^ exosomes harvested from uninfected and infected hMDMs, and OMVs purified from C-media of *P.a.* infected hMDMs to mass spectrometry analysis (LC-MS/MS). When the spectral peaks were compared between the samples several significant differences were noted (**Fig. 7C**). Next, peptide sequences were analyzed using the MASCOT database against human and *P.a.* proteomics. We identified sixty-five unique bacterial proteins in the exosomes and 79 proteins in the OMVs (**Fig. 7D**). Thirty-one bacterial proteins (**Table 2)** were present in both exosomes and OMVs isolated from C-media. The top 10 common proteins (**Fig. 7E)** are either bacterial proteases, toxins, or antigenic proteins involved in host cell modulation, suppression of immune responses, or aid in host cell and tissue damage^46–48^. These include bacterial products, such as B-type flagellin, immunomodulating metalloproteases, endopeptidase, and toxins such as exotoxin A. Thus, we found that infected host cells release both exosomes and OMVs, and most importantly the exosomes and OMVs are loaded with bacterial proteins, which are responsible for cardiomyocyte contractile dysfunction.

**Figure 7:**
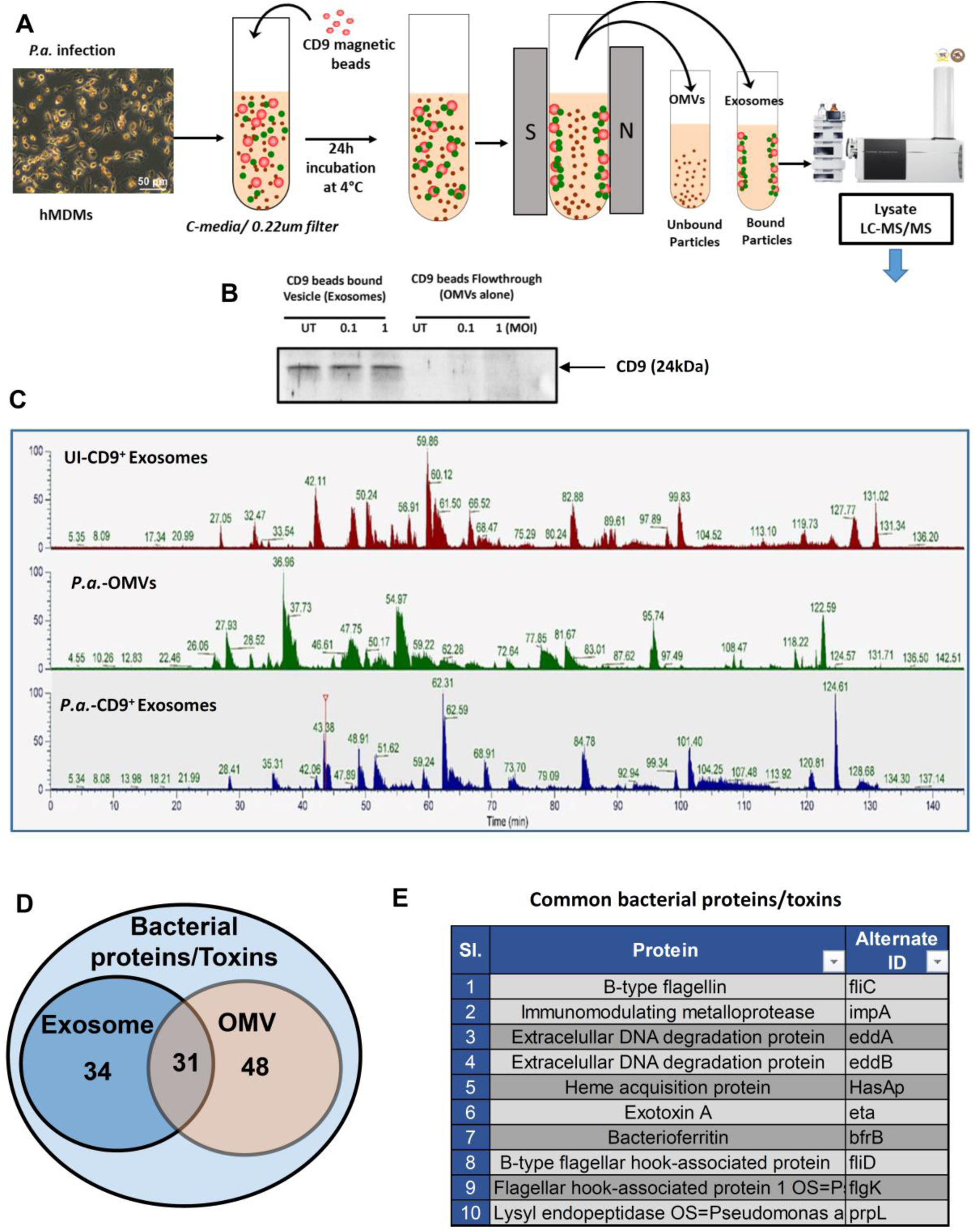
Exosomes from C-media contain bacterial antigens and products. Exosomes were harvested from uninfected and *P.a.* infected hMDMs and subject to LC- MS/MS analysis. (**A**) Schematic explains the procedure for the separation of exosomes released from macrophages, and *P.a.* derived OMVs. (**B**) Western blot analysis of lysates from exosomes (CD9 bound) and OMV (flow through) preparation, the membrane is probed with anti-CD9 antibody. (**C**) Spectral views of CD9^+^ exosomes from uninfected macrophages (top panel), purified OMVs (middle panel), and CD9^+^ exosomes from *P.a.* infected macrophages (bottom panel). (**D**) The Venn diagram shows the bacterial proteins present in CD9^+^ exosomes alone, OMVs alone, or both. (**E**) The table lists the common bacterial antigens and toxins present in either CD9^+^ exosomes or OMVs.

**Table 2:** Bacterial proteins that are common in both exosomes and OMVs. Overnight cultured hMDMs were infected with 1 MOI of *P.a.* for 22 h; the cell-free culture supernatants were used to harvest the EVs using the above-described protocol (materials and methods section). To separate the bacterial OMVs and host-derived exosomes, we incubated the EVs with antiCD9 antibody-coated beads and separated the exosomes and OMVs using magnetic columns. The purified exosomes and OMVs were subjected to LC-MS/MS analysis. The unique bacterial proteins found in both OMVs and exosomes are listed in the table.

| Sl. | Proteins | Accession Number | Alternate ID |
| --- | --- | --- | --- |
| 1 | B-type flagellin OS=Pseudomonas aeruginosa | P72151 | fliC |
| 2 | Alkaline phosphatase H OS=Pseudomonas | P35483 | phoA |
| 3 | Probable bacteriophage protein OS=Pseudomonas | Q9I5S9 | PA0623 |
| 4 | Probable bacteriophage protein OS=Pseudomonas | G3XD39 | PA0622 |
| 5 | Chaperonin GroEL OS=Pseudomonas aeruginosa | P30718 | groEL |
| 6 | B-type flagellar hook-associated protein 2 | Q9K3C5 | fliD |
| 7 | Glycerophosphoryl diester phosphodiesterase | Q9I6E6 | glpQ |
| 8 | UPF0312 protein PA0423 OS=Pseudomonas | Q9I690 | PA0423 |
| 9 | Uncharacterized protein OS=Pseudomonas | G3XD65 | PA0626 |
| 10 | Extracellular DNA degradation protein, Edd | Q9HXA3 | eddA |
| 11 | Uncharacterized protein OS=Pseudomonas | G3XCU8 | PA0628 |
| 12 | Flagellar hook-associated protein 1 OS=Pseudomonas | Q9I4P3 | flgK |
| 13 | Cytosol aminopeptidase OS=Pseudomonas | O68822 | pepA |
| 14 | Soluble pyridine nucleotide transhydrogenase | P57112 | sthA |
| 15 | Immunomodulating metalloprotease OS=Pseudomonas | Q9I5W4 | impA |
| 16 | Dihydrolipoyllysine-residue acetyltransferase | Q59638 | aceF |
| 17 | Lysyl endopeptidase OS=Pseudomonas aeruginosa | Q9HWK6 | prpL |
| 18 | Succinate--CoA ligase [ADP-forming] subunit | P53593 | sucC |
| 19 | Heme acquisition protein HasAp OS=Pseudomonas | G3XD33 | hasAp |
| 20 | Uncharacterized protein OS=Pseudomonas | Q9I0K3 | PA2635 |
| 21 | Dihydrolipoyl dehydrogenase OS=Pseudomonas | Q9I3D1 | lpdG |
| 22 | Probable bacteriophage protein OS=Pseudomonas | Q9I5S5 | PA0641 |
| 23 | Alkyl hydroperoxide reductase C OS=Pseudomonas | Q9HY81 | PA3529 |
| 24 | DNA-directed RNA polymerase subunit beta' O | Q9HWC9 | rpoC |
| 25 | Leucine-, isoleucine-, valine-, threonine-, and | P21175 | braC |
| 26 | Azurin OS=Pseudomonas aeruginosa (strain) | P00282 | azu |
| 27 | Extracellular DNA degradation protein, Edd | Q9HXA4 | eddB |
| 28 | Arginine deiminase OS=Pseudomonas aeruginosa | P13981 | arcA |
| 29 | Polyribonucleotide nucleotidyltransferase O | Q9HV59 | pnp |
| 30 | Probable TonB-dependent receptor OS=Pseudomonas | Q9HT68 | PA5505 |
| 31 | Inosine-5'-monophosphate dehydrogenase | Q9HXM5 | guaB |

To further confirm our LC-MS/MS data, lysates from CD9^+^ exosomes and CD9^-^ OMVs separated from EVs were subjected to western blot analysis and probed with anti-flagellin-B antibody. Consistent with our LC-MS/MS data we found flagellin-B in both exosomes and OMVs, which suggests that EVs contain exosomes packed with bacterial proteins and bacterial OMVs (**S. Fig. 5A**). Next, we isolated exosomes from bronchoalveolar lavage fluid (BALF) from control and *P.a.* infected mice, and lysates were examined for the presence of flagellin-B by western blot. Our data showed that exosomes from BALF of *P.a*. infected mice contain flagellin-B (**S. Fig. 5B**). Since *P.a.* infection is very common and causes serious cardiovascular complication^49,50^ in patients admitted to intensive care units (ICU), we purified exosomes from the serum of healthy donors and *P. aeruginosa* culture-positive ICU patients and determined the presence of flagellin-B. Interestingly, we found that exosomes from ICU patients contains flagellin-B at varying concentrations (**S. Fig. 5C**). These data strongly suggest that exosomes released from *P.a.* infected mice and humans carry *P.a.* antigens and toxins.

### *P.a.* OMVs cause cardiac dysfunction and increase mortality in mice

We have previously demonstrated that *P.a.* infection causes cardiac dysfunction without disseminating into the heart, and our current studies have revealed that OMVs are responsible for cardiomyocyte contractile dysfunction. To better understand the effect of OMVs on cardiac dysfunction, we performed an *in vivo* experiment by administering purified OMVs into C57BL/6J mice via the intravenous route and monitored their weight loss, survival, cardiac electrical activity, and heart function. The data showed that OMV administration caused severe mortality in mice (**Fig. 8A**), as well as arrhythmia as evidenced by irregular RR intervals and decreased heart rate (**Fig. 8B-D**). Next, we examined left ventricular functions using echocardiography. The cardiac patches shown in **Fig. 7E** clearly indicated that OMV infusion increases left ventricular wall thickness. Further analysis of cardiac patches from control and OMV infused animals revealed that OMV infusion significantly reduces stroke volume and cardiac output (**Fig. 8F-G**). Consistent with the defective left ventricular function, we found that the thickness of LV on both posterior and anterior during diastole and systole were significantly increased (**Fig. 8H-K**). As a result, the diastolic and systolic LV chamber volume was significantly less in the OMVs administrated mice (**Fig. 8L-M**). Together, these data suggest that OMVs released during infection can enter the circulation and the heart, resulting in cardiac dysfunction.

**Figure 8:**
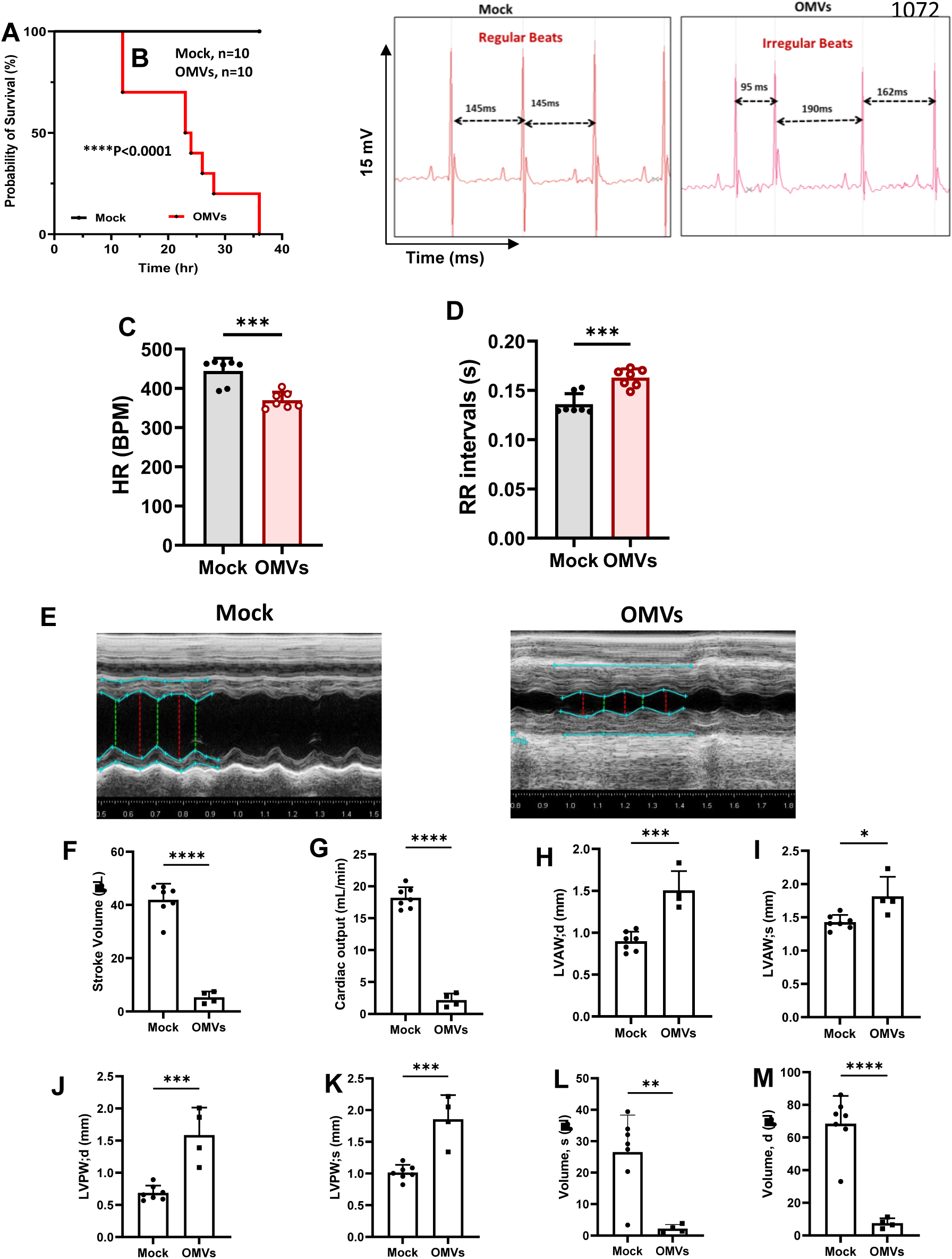
OMVs from P.a. cause cardiac electrical, LV dysfunction, and mortality in mice. C57BL/6 mice were injected intravenously with *P.a.* OMVs (10 mg/kg body weight) or saline, and survival of the mice was monitored for 36h. (**A**) The graph shows the survival rate of mice infused with OMVs (N=10). (**B**) Representative ECG traces from mock and OMV-infused mice; double head arrows indicate RR intervals. ECG traces were analyzed using Lab Chart 8 Pro (AD Instruments). The graph shown is accumulative data from 7 mice, (**C**) heart rate, and (**D**) RR intervals. To assess heart function *in vivo*, 2D- echocardiography (Vevo 2100, VisualSonics) was performed at 24h post OMVs administration. (**E**) Representative images of cardiac patches from mock and OMVs administrated mice. At least 3 M-mode echocardiogram measurements from each mouse were used to determine the (**F**) stroke volume, (**G**) cardiac output, (**H-I**) anterior, (**J-K**) posterior LV wall thickness, and (**L-M**) LV chamber volume during diastole and systole. The data shown are representative of 2 different experiments (Mock, n=7, and OMVs, n=4), mean ± SD **p<0.01; **p<0.001; ***p<0.0001.

## Discussion

Pneumonia is a leading cause of morbidity and mortality in the world^51,52^. Bacterial infections such as *Streptococcus pneumoniae* and viral infections such as influenza or SARS-CoV-2 are the most common causes of pneumonia^7,53–55^. Irrespective of the infecting agent, the comorbidity of pneumonia and its associated cardiovascular complication increases the mortality rate in those admitted to the ICU^7,53,56,57^. We showed that intranasal instillation of *P.a.* increases mortality and causes left ventricular (LV) dysfunction despite limited dissemination of bacteria into the heart^10^. This suggests that other mediators released from the host and/or pathogen play a vital role in aggravating LV dysfunction during *P.a.* infection. However, the mechanism of how *P.a.* lung infections drive cardiovascular complications, and their associated mortality is unclear. In the present work, using *P.a.* model organism for pneumonia and we investigated the mechanism of pneumonia-induced cardiac dysfunction.

The nature and potential function of C-media depend on the cell types, source, and pre-existing state. For example, the C-media from amniotic progenitor cells and mesenchymal stem cells have antimicrobial and antitumor properties^58–60^. Conversely, when the source and environment change, C-media can contribute to adipocyte inflammation, cancer metastasis, and the release of inflammatory cytokines^61,62^. In the present study, we found that cell-free C-media harvested from *P.a.* infected hMDMs cause cardiomyocyte contractile dysfunction. *P.a.* C-media decreases the beat period, and consequently, there was tachycardia in treated cardiomyocytes as early as 45 min of exposure. After 2h, the contraction of cardiomyocytes was eliminated (**Fig. 1**). However, the morphology and survival of treated cardiomyocytes were intact, and we found 45-55% cell death at 24h post-treatment (**S.Fig. 1A-B).** Furthermore, reduced FPD showed that depolarization and repolarization occurred for a shorter duration in treated cardiomyocytes (**Fig. 1D**). If we compare this result in the context of the actual heart, this phenomenon could reduce cardiac output by reducing diastolic filling time and ventricular filling^63^ as we reported in our earlier publication^10^.

Intercellular gap junction proteins and connexin 43 help to connect cardiomyocytes with neighboring cells; this process is critical for establishing synchronized wave propagation and electrical conduction^64,65^. The uniform beat period and normal electrical conduction wave in the cardiomyocytes treated with C-media from uninfected hMDMs confirm that these cardiomyocytes were well synchronized. However, the slower and prolonged electrical conduction wave in *P.a.* C-media-treated cardiomyocytes indicates malfunction of electrical conduction and damage in the syncytium of the cardiomyocytes. The systemic influx and efflux of ions tightly regulate cardiac action potentials. The rapid depolarization of membrane potential, driven by Na^+^ influx, initiates the action potential’s upstroke. A closure examination of MEA data reveals phase 2 EADs of the action potential in the hiPSC-CMs exposed to *P.a.* C-media. This phenomenon resembles the Na^+^ channel inactivation mechanism observed with ATX-II treatment, a peptide toxin derived from sea anemone and known to inhibit the Na^+^ channel function^66,67^. Similarly, the enhanced entry of Na^+^ through the sarcolemma into the cytoplasm triggers a concomitant rise in intracellular Ca^2+^, a key factor known to induce arrhythmias^43,68^. Our findings revealed that in hiPSC-CMs treated with *P.a.* C-media, intracellular Ca^2+^ levels were elevated, along with irregular Ca^2+^ cycling intervals. This irregularity in Ca^2+^ handling suggests a disruption in cardiac excitation-contraction coupling, leading to arrhythmogenesis. Importantly, this study is the first to demonstrate that OMVs from *P.a.* can modulate Na^+^ currents, thereby impacting both Na^+^ and Ca^2+^ homeostasis and contributing to arrhythmogenic potential.

In response to bacteria PAMPs, the host cell produces inflammatory cytokines such as IL-1β, IL-6, and TNF-α to activate innate immune cells including macrophages, monocytes, and neutrophils^69–72^. Recent studies revealed that host cells can release EVs, that are packed with various particulates such as cytokines, small RNAs, microRNAs, etc., which can initiate an inflammatory response^73^. Furthermore, bacteria also release a very high number of OMVs while residing in the host cells or in the extracellular environment^40,72^. In this study, we found that *P.a.* infected macrophages released significantly elevated levels of cytokines and EVs that are packed with bacterial OMVs, which suggests that these cytokines and EVs could be potent mediators of cardiomyocyte contractile dysfunction. To validate whether the cardiomyocyte contractile dysfunction is caused by inflammatory cytokines, our experiments demonstrated that heat treatment of C-media to inactivate cytokines actually aggravated the cardiomyocyte contractile dysfunction. This observation would seem to rule out the involvement of heat-sensitive cytokines^74,75^ in cardiomyocyte contractile dysfunction. We interpret this result to indicate that heat inactivation of C-media leads to EVs membrane rupture and release of the bacterial proteins contents that mediate the cardiomyocyte contractile dysfunction. This interpretation was further supported by our data obtained from the hiPSC-CMs exposed to C-media harvested from hMDMs treated with heat-killed *P.a.* or live *P.*a., which showed that only C-media harvested from hMDMs treated with live *P.a.* mediated cardiomyocyte contractile dysfunction.

To find out more about the involvement of DAMPs and PAMPs on cardiomyocyte contractile dysfunction, *P.a.* were infected in culture medium with or without macrophages. Our data revealed that both media from *P.a.* cultures, which contains secreted PAMPs and OMV, and C-media from macrophages containing EVs packed with both DAMPs and PAMPs showed identical cardiomyocyte contractile dysfunction by decreasing the beat period. Thus, cardiomyocyte contractile dysfunction is primarily due to PAMPs released from infected macrophages in exosomes and OMVs.

Bacteria use OMVs to communicate with host cells and modulate the host immune system to create a niche for colonization, transmission of virulence factors, and establishment of pathogenesis^19,76^. Herein, we demonstrated that OMVs produced by *P.a.* cause cardiomyocyte contractile dysfunction as evidenced by increasing the beat period and subsequent decrease in the beat rate. Furthermore, the irregular beat period and missing depolarization spike of cardiomyocytes confirmed that OMV treatment also induces arrhythmias. Since the induction of arrhythmias is directly associated with Ca^2+^ cycling disturbance^43,77,78^, we examined Ca^2+^ handling by the cardiomyocyte during *P.a.* C-media and OMV treatment. Importantly, both *P.a*. C-media and OMVs altered the intracellular Ca^2+^ oscillation of the cardiomyocytes by changing t parameters such as peak amplitude, peak duration, time-to-peak ratio, and decay time.

Since Infected macrophages release EVs and bacteria release OMVs into the medium^17,40^, we proceeded to determine the impact of the contents of exosomes or OMVs from *P.a.* infected hMDMs on cardiomyocytes. An important question was whether the exosomes produced by macrophages during *P.a.* infection carry molecules of bacterial origin. Our results support this hypothesis, since exosomes produced by hMDMs during infection contained bacterial proteins. This was demonstrated by our LC-MS/MS data which revealed that thirty-one bacterial proteins/toxins identified in OMVs were also detected in exosomes isolated from the *P.a.* C-media. Hence, this confirmed that exosomes generated during bacterial infection carry bacterial-originated biomolecules. To validate the LC-MS/MS finding, we isolated exosomes from BALF of *P.a.* infected mice and the serum of ICU patients positive for *P.a.* infection and found that flagellin-B was loaded in these exosomes. These findings further supported our LC-MS/MS data and confirmed that exosomes released during bacterial infection carry biomolecules from *P.a*.

Our *in-vitro P.a.* infection model revealed that exosomes released from infected macrophages and OMVs released from *P.a.* carry virulent factors that can damage organs, including the heart. Furthermore, we found that OMVs intravenously administered to C57BL/6 mice caused significant cardiac abnormalities (LV wall thickness, less cardiac output, etc.) and increased mortality.

In summary, we demonstrated that C-media from *P.a.* infected hMDMs cause cardiomyocyte contractile dysfunction. Our data defines the role of exosomes and OMVs on ion channels and their downstream effect on cardiomyocyte contractile function. The mechanism that we found in our *in vitro* cell culture model with hiPSC-CMs was recapitulated in our *in vivo* model, in which when we delivered the bacterial OMVs into the systemic circulation, it caused severe cardiac abnormalities, cardiac arrests, and increased mortality. Thus, neutralizing the major bacterial antigens packed in OMVs could be potential therapeutics for patients admitted to ICU with *P.a.* pulmonary infections.

## Acknowledgment

The authors thank The Ohio State University Campus Chemical Instrument Center (CCIC) Mass Spectrometry (The Fusion Orbitrap instrument was supported by NIH Award Number Grant S10 OD018056), Animal care and use program, and Small Animal Imaging Core Lab for all our imaging studies. Special Thanks to Dr. Sakthivel Sadayappan, Ph.D, Chiar, Department of Cellular and Molecular Medicine, University of Arizona, Tucson, AZ for editing the manuscript.

## Authors Contribution

N.K., D.J.W. and M.V.S.R designed the research; N.K., S.S.M., S.S, M.P.E, S.M., CP., W.P.L., and M.V.S.R. performed the research; N.K., S.S.M., S.S, W.P.L., D.J.W., L.P.G. H.S, M.K., and M.V.S.R analyzed the data; J.B. provided human samples. N.K., S.S.M, and M.V.S.R. wrote the manuscript, and S.S., L.P.G., W.P.L., H.S., D.J.W and M.V.S.R edited the manuscript.

---

**Supplemental Figure 1:**
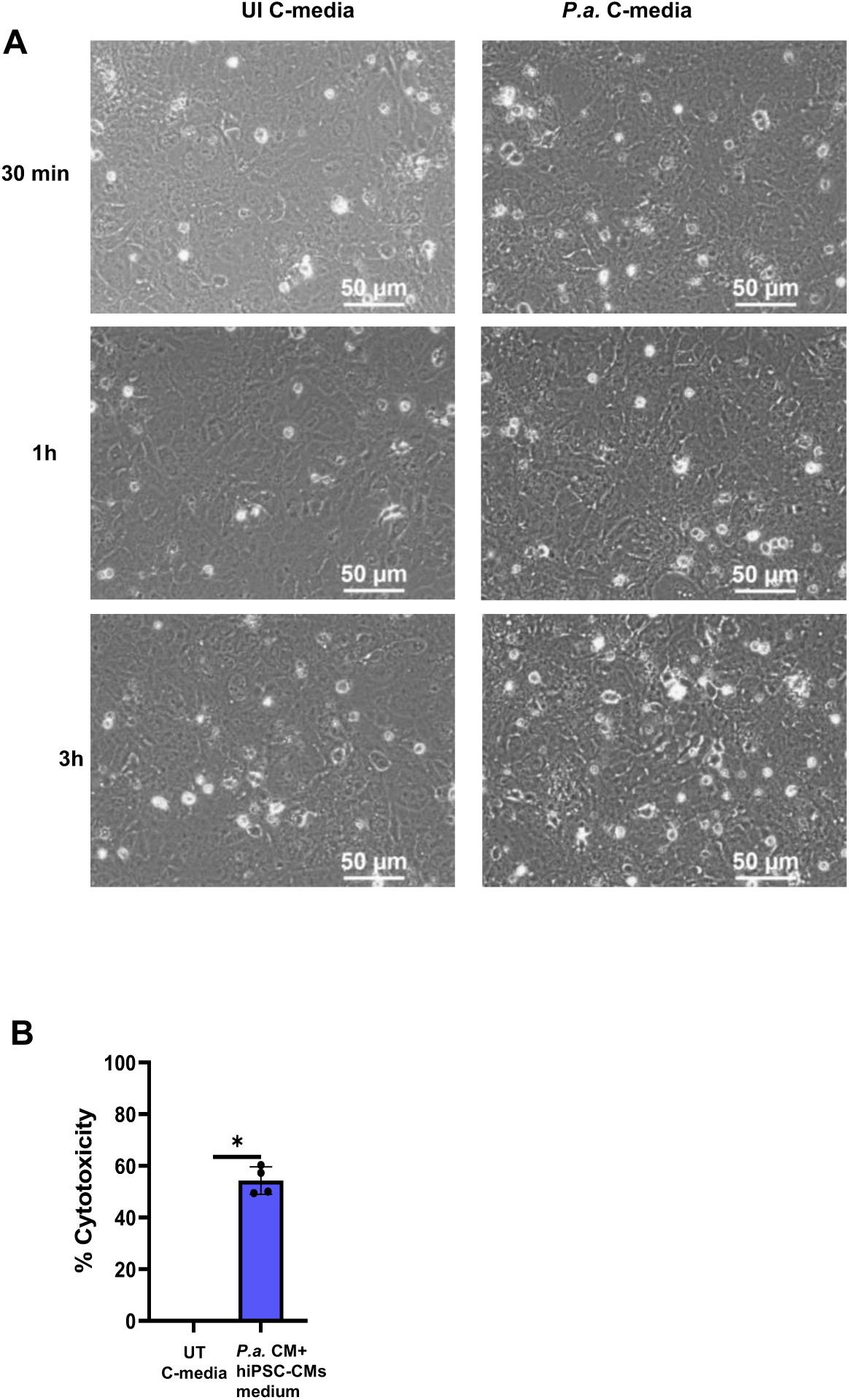
Cytotoxicity of cardiomyocytes treated with C-media from ***P.a.* infected hMDMs.** hiPSC-CMs were plated in 12 well-sterile, and the cells were exposed to the mixture of cardiomyocyte culture medium and C-media (1:1 ratio) harvested from uninfected (UI C-media) or *P.a.* infected (*P.a.* C-media) hMDMs. (**A**) Representative Phase-contrast image of hiPSC-CMs cultured for 7-10 days before the exposure to C-medium. Images were captured at 30 min, 1h, and 3h post-exposure to UI C-media and *P.a.* C-medium. Images were acquired on an Olympus CKX41 Phase contrast microscope fitted with Leica MC170 HD. Scalebar, 50µm. The images shown are representative of three independent experiments. (**B**) Cell free culture supernatants were harvested after 24 hours of exposure of hiPSC-CMs and used for cytotoxicity assay (LDH assay). Graph shown is representative of three independents experiments; mean ± SD; * p<0.05.

**Supplemental Figure 2:**
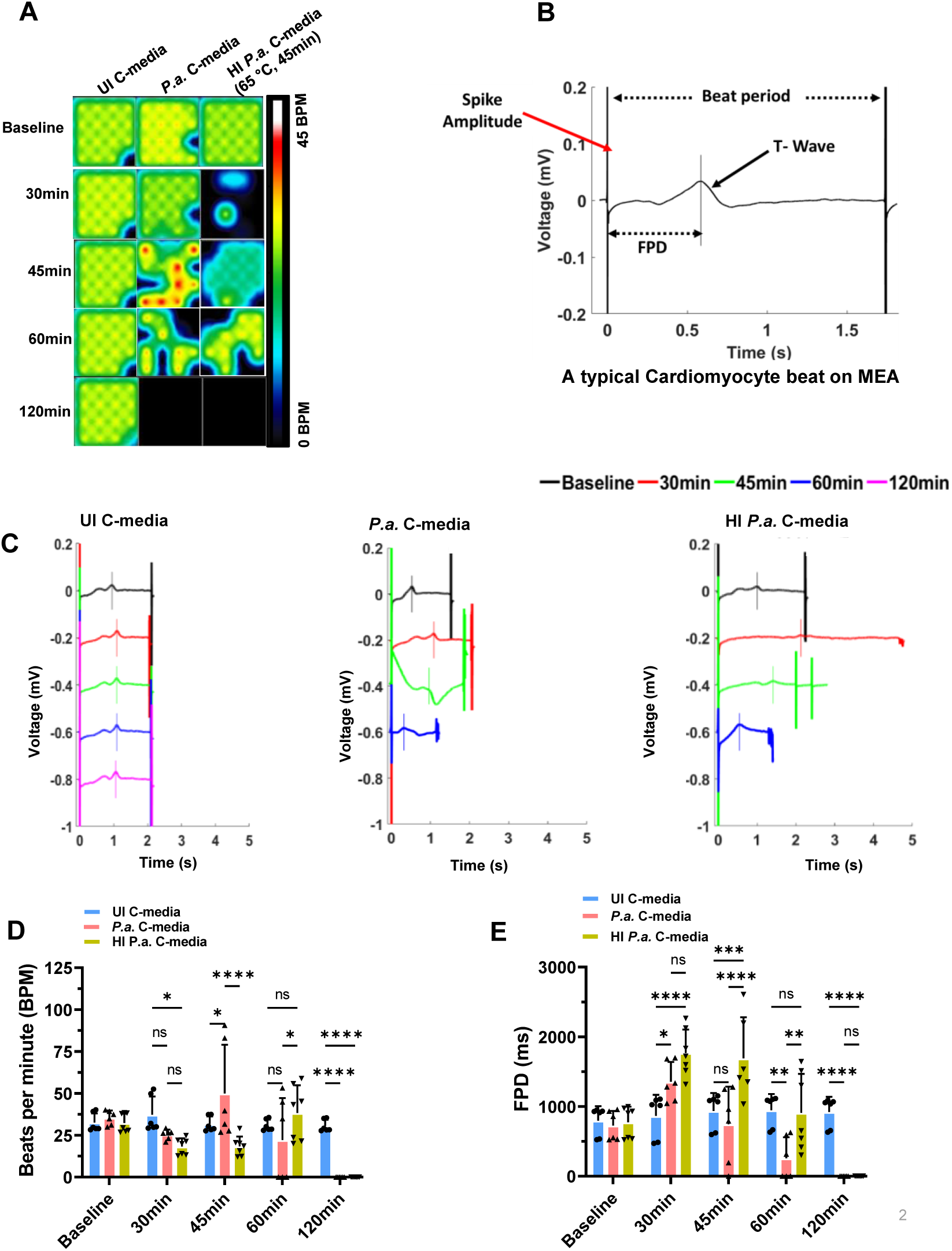
Heat-inactivated C-media from P.a. infected hMDMs causes cardiomyocyte contractile dysfunction. hiPSC-CMs were plated in 24 well MEA plates, and the cells were exposed to the mixture of cardiomyocyte culture medium and C-media (1:1 ratio) harvested from hMDMs that were left uninfected (UI C-media) or P.a. infected (P.a. C-media) and heat-inactivated C-media (at 65°C for 45 min). The cardiomyocyte contractility and electrical activity was recorded using AxIS Navigator on MEA system at 5% CO2 and 37°C for indicated time points. Data analysis was performed using the cardiac analysis tool. (**A**) Electrical activity map showing the changes in the beat rate., Shown is a representative well from quadruplicate wells from each treatment (N=3). (**B**) A typical Cardiomyocyte beat on MEA. (**C**) Representative cardiomyocyte beat overlay recorded with the MEA system showing the beat period, T-wave, and FPD in UI C-media, P.a. C-media, and heat-inactivated C-media treated hiPSC-CMs. The graphs shown in (**D**) are beat rate, and (**E**) are field potential duration (FPD) at baseline, 30-, 60-, and 120 minutes post-treatment of hiPSC-CMs. Data shown in figures D and E are accumulative data from 3 independent experiments. mean ± SD; * p<0.05, ** p <0.01, *** p < 0.001, **** p < 0.0001.

**Supplemental Figure 3:**
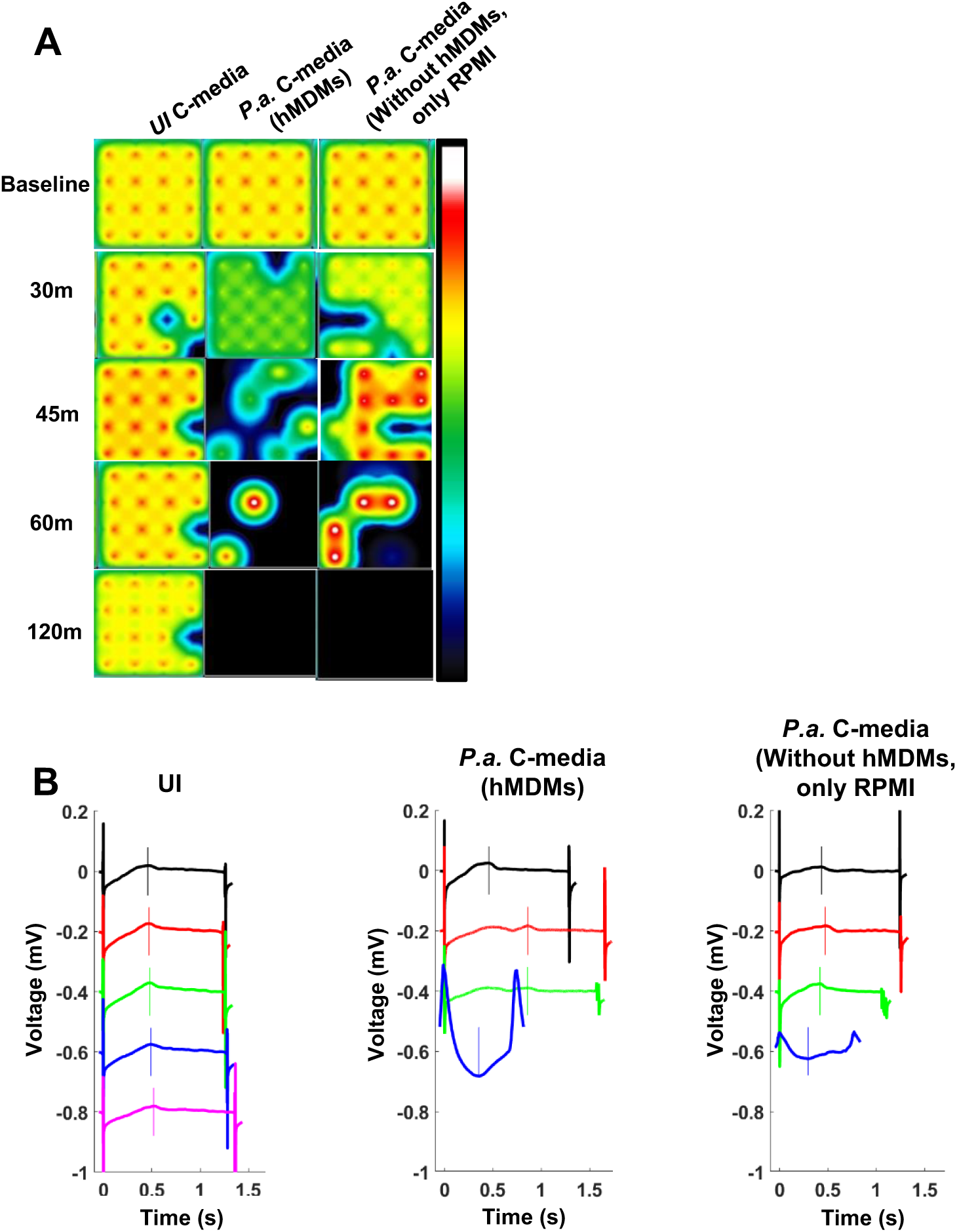
Direct infection of *P.a.* delays cardiomyocyte contractile dysfunction. hiPSC-CMs were plated in 24 well MEA plates, and the cells were exposed to *P.a.* (1 MOI) or left uninfected (UI). The cardiomyocyte contractility and electrical activity was recorded using AxIS Navigator on the MEA system at 5% CO2 and at 37°C for indicated time points. Data analysis was performed using the cardiac analysis tool. (**A**) Electrical activity map showing the changes in the beat rate of cardiomyocytes infected with *P.a.* or control (UI). The activity map shown is a representative well from quadruplicate samples from each treatment (N=3). (**B**) A representative cardiomyocyte beat overlay was recorded with the MEA system, showing the beat period, T-wave, and FPD in the UI and *P.a.* infected hiPSC-CMs. The graphs shown in (**C**) are beat rate, and (**D**) are field potential duration (FPD) at baseline, 5h, 6h, 7h, and 8h post-infection of hiPSC-CMs with *P.a*. or UI. Data shown in figures D, E, and F are accumulative data from 3 independent experiments; mean ± SD; * p<0.05, ** p <0.01, *** p < 0.001, **** p < 0.001, *** p < 0.0001. (**G**) Representative Phase-contrast images were captured at 8h post-infection with *P.a.* or left uninfected. Scalebar=50µm. The images shown are representative of three independent experiments.

**Supplemental Figure 4:**
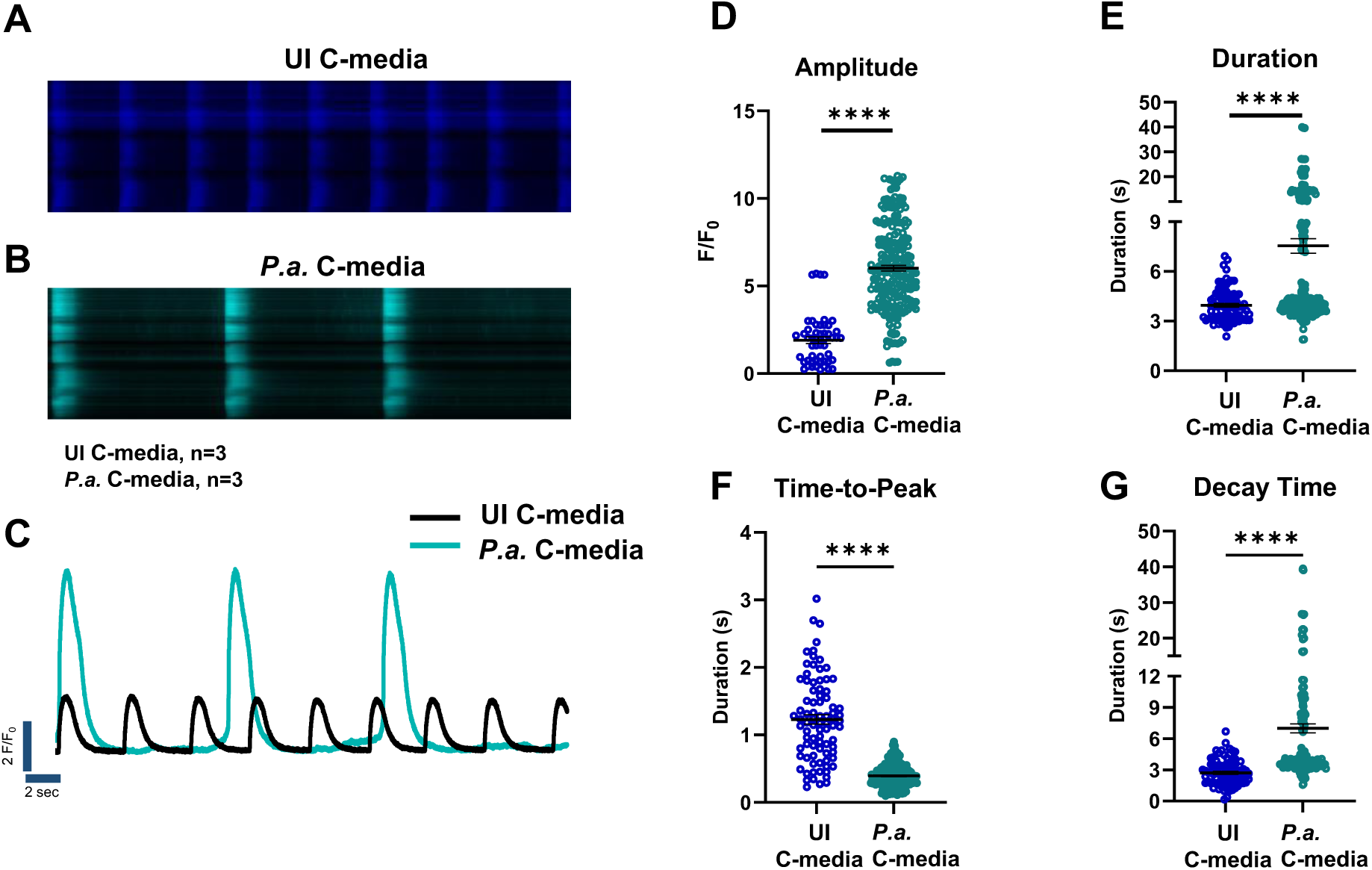
C-media from bacterial culture causes cardiomyocyte contractile dysfunction. *P.a.* C-medium was collected as described earlier. For culturing *P.a.* without hMDMs, *P.a.* was grown in RPMI supplemented with 10% autologous serum (serum used for hMDM culture) for 2 days, and C-media was processed as described earlier. hiPSC-CMs were plated in 12 well-sterile plates, and the cells were exposed to the mixture of cardiomyocyte culture medium and C-media (1:1 ratio) harvested from uninfected (UI C-media), *P.a.* infected (*P.a.* C-media) hMDMs or *P.a* RPMI C-media without hMDMs. The cardiomyocyte contractility and electrical activity were recorded using AxIS Navigator on the MEA system at 5% CO2 and at 37°C for indicated time points. Data analysis was performed using the cardiac analysis tool. (**A**) Electrical activity map showing the changes in the beat rate of cardiomyocytes infected with *P.a.* C-media with/without hMDMs or control (UI). The activity map shown is a representative well from quadruplicate samples from each treatment (N=3). (**B**) A representative cardiomyocyte beat overlay was recorded with the MEA system, showing the beat period, T-wave, and FPD from different experimental conditions as described above.

**Supplemental Figure 5:**
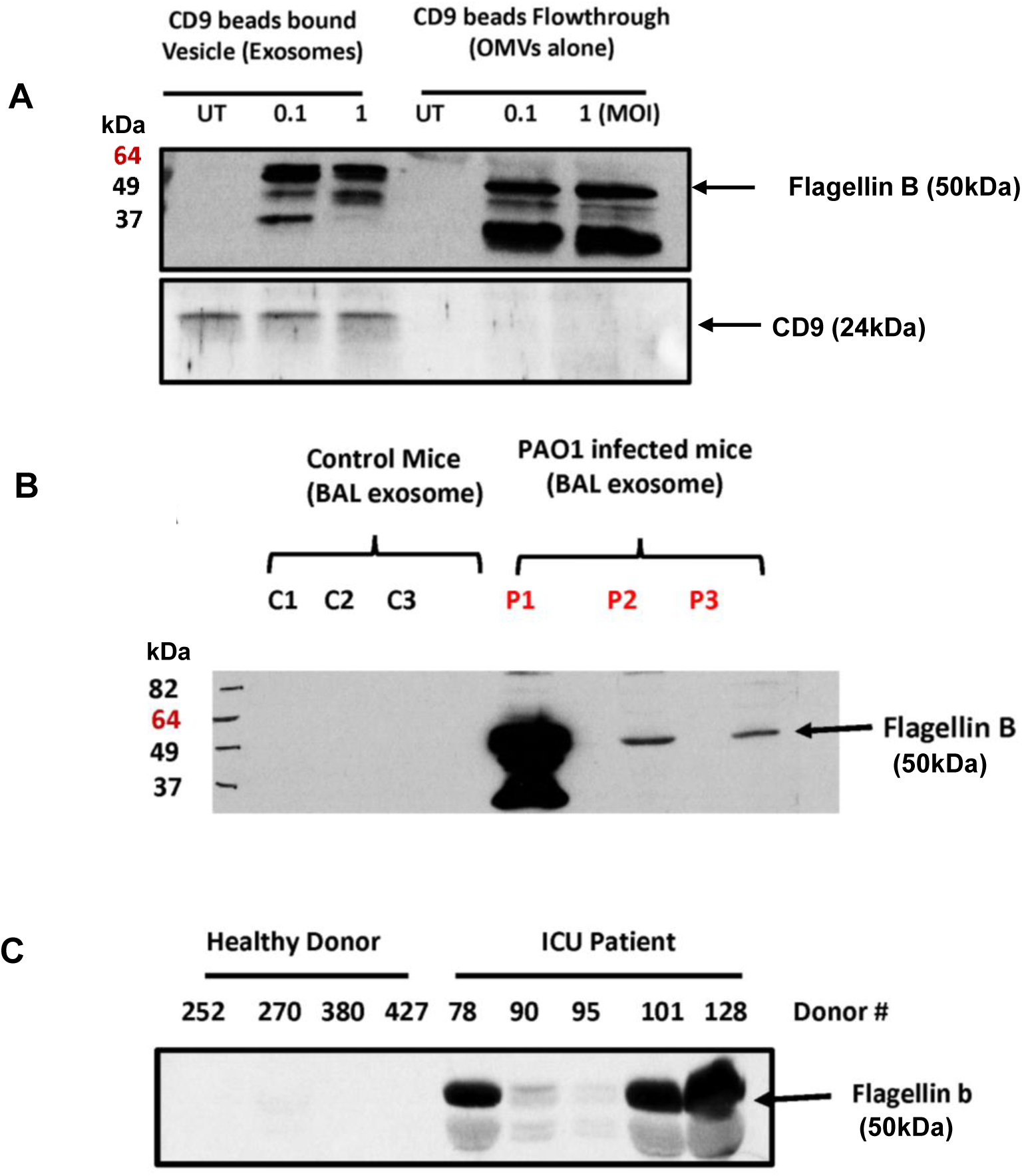
***P.a.* Flagellin-B is loaded in exosomes isolated from hMDMs, mouse BALF and serum from ICU patients**. Western blot analysis shows that exosomes were carry flagellin-B. (**A**) Western blot analysis of lysates from exosome and OMVs separated from *P.a.* C-medium using CD9 beads, (**B**) exosomes from BALF of *P.a.* infected C57BL/6J mice and (**C**) exosomes from human serum from healthy donors and ICU patients positive for *P.a.* infection. The membrane was probed with anti-flagellin- B antibody and CD-9 antibody.

